# ID transcription factors regulate the ability of Müller glia to become proliferating neurogenic progenitor-like cells

**DOI:** 10.1101/2023.10.02.560518

**Authors:** Olivia B. Taylor, Snehal P. Patel, Evan C. Hawthorn, Heithem M. El-Hodiri, Andy J. Fischer

## Abstract

The purpose of this study was to investigate how ID transcription factors (TFs) regulate the ability of Müller glia (MG) to reprogram into proliferating MG-derived progenitor cells (MGPCs) in the chick retina. We found that *ID1* is transiently expressed by maturing MG, whereas *ID4* is upregulated and maintained in maturing MG in embryonic retinas. In mature retinas, *ID4* was prominently expressed by resting MG, but in response to retinal damage *ID4* was rapidly upregulated and then downregulated in MGPCs. By contrast, *ID1, ID2* and *ID3* were low in resting MG and then upregulated by MGPCs. Inhibition of ID TFs following retinal damage decreased numbers of proliferating MGPCs. Inhibition of IDs after the proliferation of MGPCs significantly increased numbers of progeny that differentiate as neurons. In damaged or undamaged retinas inhibition of IDs increased levels of p21^Cip1^ in MG. In response to damage or insulin+FGF2 levels of *CDKN1A* message and p21^Cip1^ protein were decreased, absent in proliferating MGPCs, and elevated in MG returning to a resting phenotype. Inhibition of Notch- or gp130/Jak/Stat-signaling in damaged retinas increased levels of ID4 but not p21^Cip1^ in MG. Although *ID4* is the predominant isoform expressed by MG in the chick retina, *id1* and *id2a* are predominantly expressed by resting MG and downregulated in activated MG and MGPCs in zebrafish retinas. We conclude that ID TFs have a significant impact on regulating the responses of MG to retinal damage, controlling the ability of MG to proliferate by regulating levels of p21^Cip1^, and suppressing the neurogenic potential of MGPCs.

## Introduction

There is a growing body of work describing how retinal Müller glia (MG) can be reprogrammed into MG-derived progenitor cells (MGPCs). MG are the predominant type of support cell in the retina and are common to all vertebrate classes. MG provide many important functions in the retina including structural, synaptic, osmotic homeostasis, and nutritive/metabolic support. However, MG also have the extraordinary capability to de-differentiate, proliferate, acquire progenitor phenotype, and generate new neurons (Fischer and Reh, 2001; Fausett and Goldman, 2006; Bernardos et al., 2007; Karl et al., 2008). Cellular reprogramming underlies the transition of MG into progenitor cells and this process involves the downregulation of glial genes, up-regulation of genes associated with progenitor cells and proliferation, followed by neuronal differentiation. In the chick retina, neuronal damage stimulates numerous MG to de-differentiate, re-enter the cell-cycle, and express transcription factors (TFs) associated with embryonic retinal progenitors (Fischer and Reh, 2001). The majority of cells generated by MGPCs remain as un-differentiated progenitor cells, some differentiate into MG and a few differentiate into neurons (Fischer and Reh, 2001; Ghai et al., 2010; Todd et al., 2016a, 2018). MG in the fish retina have an extraordinary capacity to regenerate retinal neurons whereas the capacity of MG in mammals is very poor (reviewed by (Fischer, 2005; Barbosa-Sabanero et al., 2012; Gallina et al., 2014a; Wan and Goldman, 2016; Todd and Reh, 2022). These observations raise important questions as to why fish MG regenerate numerous functional neurons, whereas this regenerative potential is restricted in birds and blocked in mammals.

We have recently reported that, unlike MG in birds and fish, mammalian MG active networks of genes to suppress reprogramming and restore glial quiescence (Hoang et al., 2020). This study implicated different pro-quiescence TFs and cell signaling pathways that actively suppress reprogramming of MG into neurogenic MGPCs. For example, in adult damaged mouse retinas, MG can be reprogrammed into neurons by knock-out the pro-glial TFs *Nfia, Nfib* and *Nfix* (Hoang et al., 2020). Alternatively, rodent MG can be reprogrammed into functional neuron-like cells by the combination of damage, HDAC inhibition and forced expression of the pro-neural transcription factor *Ascl1* (Jorstad et al., 2017); this reprogramming can be significantly increased by the ablation of microglia (Todd et al., 2020) or pharmacological inhibition of Jak/Stat (Jorstad et al., 2020), NFkB, TGFβ/Smad3 or ID TFs (Palazzo et al., 2022). Interestingly, combined forced expression of *Ascl1* and *Atoh1* in MG effectively drives reprogramming into neuronal cells in adult mice without the need for neuronal damage, even though *Atoh1* is not normally expressed in the retina (Todd et al., 2021).

The purpose of this study is to investigate how ID TFs influence the proliferation and neurogenic potential of MGPCs in the chick model system. ID proteins belong to class V of the helix-loop-helix (HLH) transcription factor family. Unlike other HLH proteins, IDs lack a basic region; instead of binding to DNA, they heterodimerize and repress the activity of class I and II bHLH proteins (reviewed by (Torres-Machorro, 2021). These TFs function to regulate differentiation, promote cell-cycle progression, senescence, angiogenesis, tumorigenesis, and metastasis in cancer (reviewed by (Murugesan et al., 2023). In the developing nervous system, ID proteins prevent premature differentiation and maintain the proliferation of progenitors via antagonism of proneural bHLH TFs and Retinoblastoma tumor suppressor-related proteins (reviewed by (Iavarone and Lasorella, 2004). The transition of progenitors from proliferation to neurogenesis involves a coordinated increase of proneural bHLH factors and a decrease in Hes and ID factors. As development proceeds, inhibition of proneural bHLH factors promotes the formation of astrocytes (reviewed by (Ross et al., 2003). Herein, we investigate patterns of expression of ID factors in retinal cells and how inhibition of IDs influences the proliferation and neurogenic potential of MGPCs in the chick model system.

## Methods

### Animals

The animals approved for use in these experiments were in accordance with the guidelines established by the National Institutes of Health and IACUC at The Ohio State University. Newly hatched P0 wildtype leghorn chicks (Gallus gallus domesticus) were obtained from Meyer Hatchery (Polk, Ohio). Post-hatch chicks were maintained in a regular diurnal cycle of 12 hours light, 12 hours dark (lights on at 8:00 AM). Chicks were housed in stainless-steel brooders at 25°C and received water and Purina^tm^ chick starter ad libitum.

### Intraocular injections

Chicks were anesthetized with 2.5% isoflurane mixed with oxygen from a non-rebreathing vaporizer. The technical procedures for intraocular injections were performed as previously described (Fischer et al., 1998). With all injection paradigms, both pharmacological and vehicle treatments were administered to the right and left eye respectively. Compounds were injected in 20 µl sterile saline. Compounds included: NMDA (38.5nmol or 154 µg/dose; Sigma-Aldrich), FGF2 (250 ng/dose; R&D systems), and insulin (1 µg/dose; Sigma-Aldrich). 5-Ethynyl-2’-deoxyuridine (EdU; 2.3 µg/dose) was intravitreally injected to label the nuclei of proliferating cells. Injection paradigms are included in each figure.

**Table 1.**
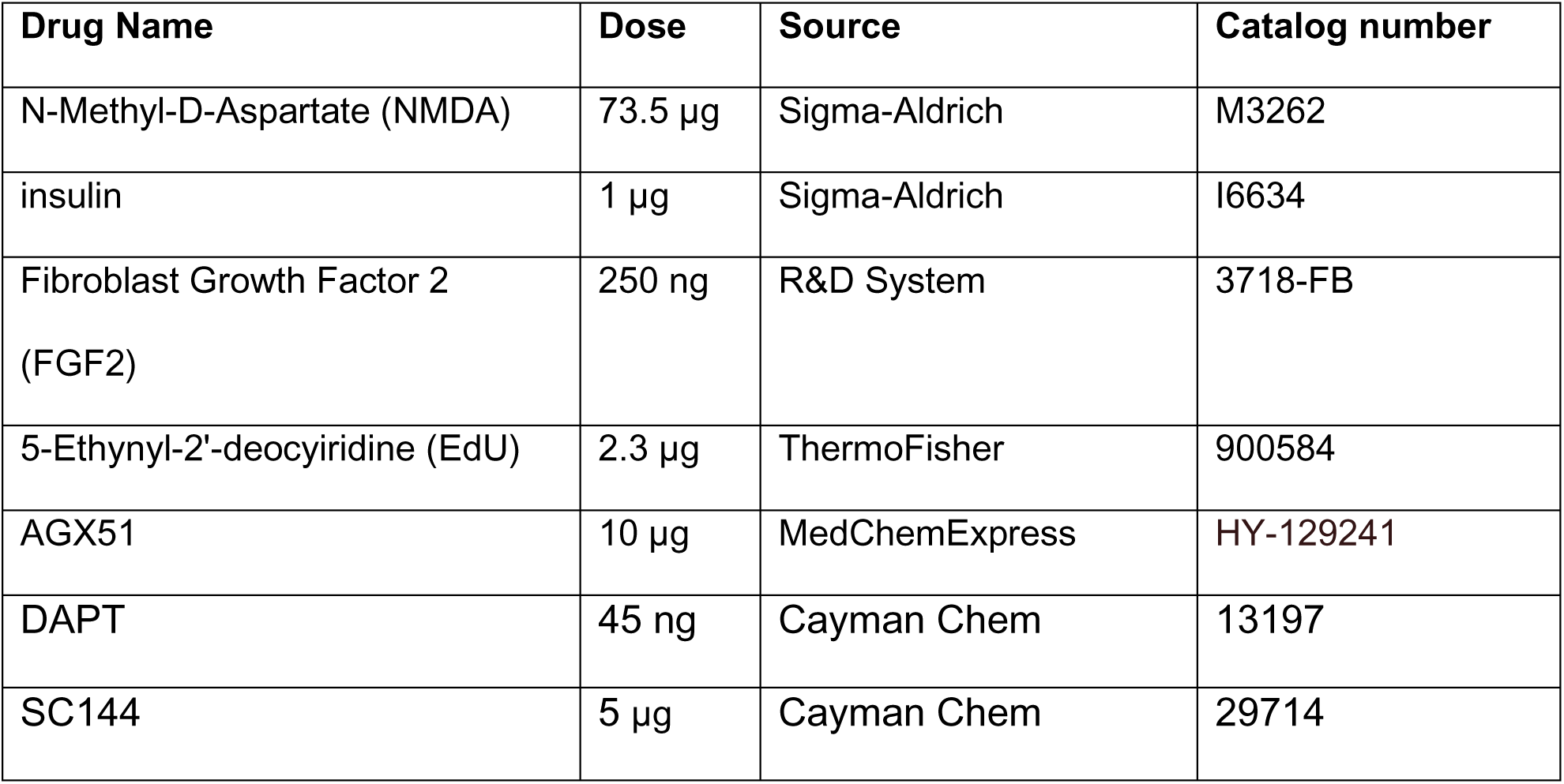
Pharmacological Compounds.

| Drug Name | Dose | Source | Catalog number |
| --- | --- | --- | --- |
| N-Methyl-D-Aspartate (NMDA) | 73.5 μg | Sigma-Aldrich | M3262 |
| insulin | 1 μg | Sigma-Aldrich | I6634 |
| Fibroblast Growth Factor 2 (FGF2) | 250 ng | R&D System | 3718-FB |
| 5-Ethynyl-2'-deocytiridine (EdU) | 2.3 μg | ThermoFisher | 900584 |
| AGX51 | 10 μg | MedChemExpress | HY-129241 |
| DAPT | 45 ng | Cayman Chem | 13197 |
| SC144 | 5 μg | Cayman Chem | 29714 |

### Fixation, sectioning, and immunocytochemistry

Retinas were fixed, sectioned, and immunolabeled as described previously (Fischer et al., 2008, 2014a; Gallina et al., 2015a). To identify MG and astrocytes we labeled sections for Sox2, Sox9, or S100β. Antibodies to Sox2 and Sox9 are known to label the nuclei or MG in the INL and the nuclei of astrocytes at the vitread surface of the retina, whereas antibodies to S100β label the cytoplasm of astrocytes (Fischer et al., 2010). None of the observed labeling was due to non-specific labeling of secondary antibodies or auto-fluorescence because sections labeled with secondary antibodies alone were devoid of fluorescence. Primary antibodies used in this study are described in **Table 2**. Secondary antibodies included donkey-anti-goat-Alexa488/568 (Life Technologies A3214; A3214), goat-anti-rabbit-Alexa488/568 (Life Technologies A32731; A-11036); and goat-anti-mouse-Alexa488/568/647 (Life Technologies A32723; A-11004; A32728) diluted to 1:1000 in PBS plus 0.2% Triton X-100.

**Table 2:**
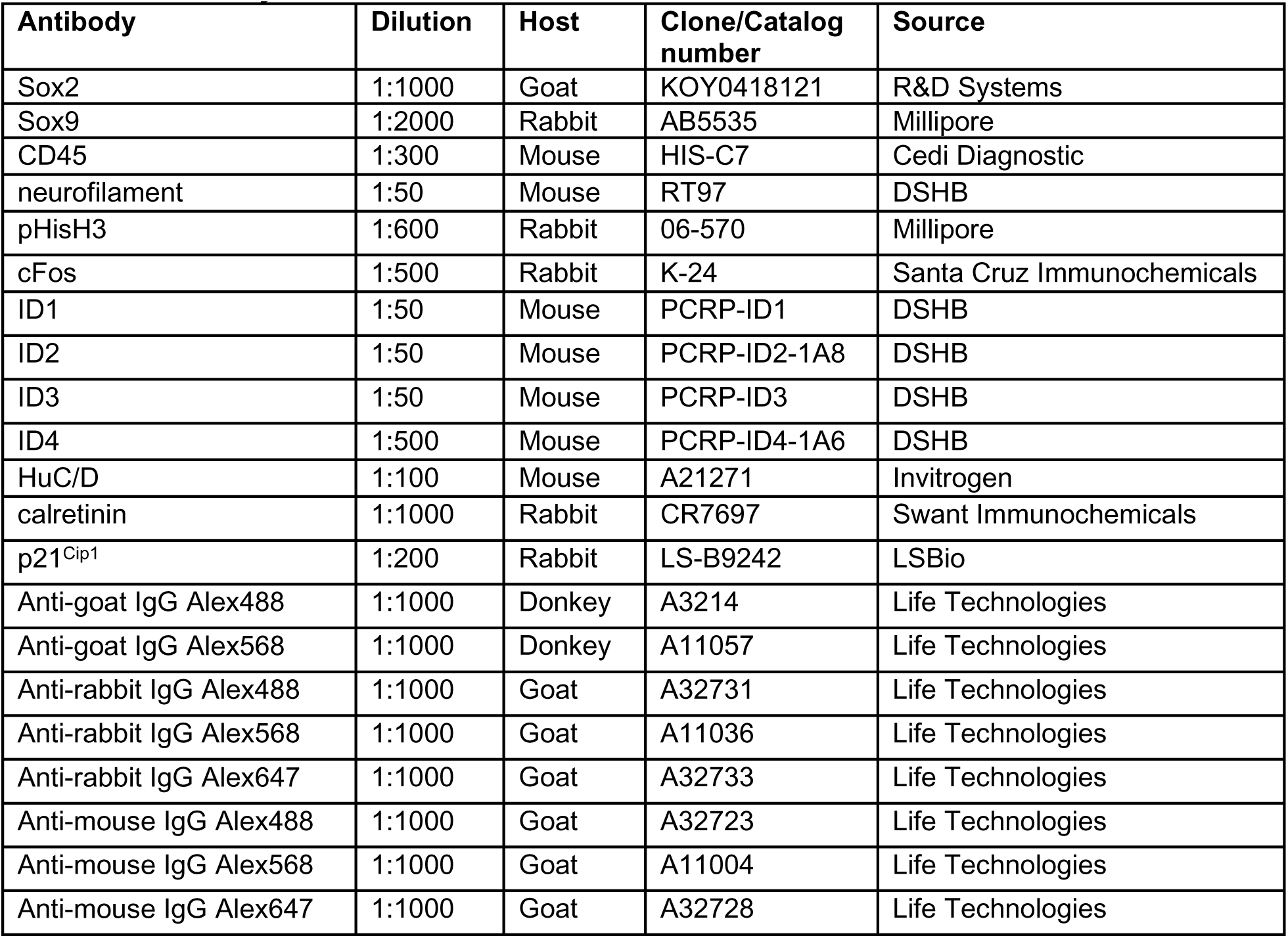
Antibody table.

### Terminal deoxynucleotidyl transferase dUTP nick end labeling (TUNEL)

The TUNEL assay was implemented to identify dying cells by imaging fluorescent labeling of double stranded DNA breaks in nuclei. The *In Situ* Cell Death Kit (TMR red; Roche Applied Science) was applied to fixed retinal sections as per the manufacturer’s instructions.

### scRNA-seq and scATAC-seq

In short, retinas were acutely dissociated via papain digestion (Worthington Biochemicals). Dissociated cells were loaded onto the 10X Chromium Cell Controller with Chromium 3’ V2 or V3 reagents. Sequencing and library preparation was performed as previously described (Hoang et al., 2020; Palazzo et al., 2020; Campbell et al., 2021b; c). Cell Ranger output files for Gene-Cell matrices for scRNA-seq data for libraries from saline and NMDA-treated retinas are available through GitHub: https://github.com/jiewwwang/Singlecell-retinal-regeneration or Sharepoint: [chick embryonic retina scRNA-seq Cell Ranger outs](https://osumc.sharepoint.com/:f:/s/Links/EtBjFrvATk9AuO_UBNr_LQABWnxJLKcLdfeVMTs3bPzjzw?e=FU3Sll) and [chick retina scRNA-seq Cell Ranger output files](https://osumc.sharepoint.com/:f:/s/Links/Eoto-Qg2uuxDn1bHWMM6gdkBTft4S_YSBjQJResxY-qehA?e=xMKHgS). scRNA-seq datasets are deposited in GEO (GSE135406, GSE242796) and Gene-Cell matrices for scRNA-seq data for libraries from saline and NMDA-treated retinas are available through NCBI (GSM7770646, GSM7770647, GSM7770648, GSM7770649).

Using Seurat toolkits (Powers and Satija, 2015; Satija et al., 2015), Uniform Manifold Approximation and Projection (UMAP) for dimensional reduction plots were generated from 9 separate cDNA libraries, including 2 replicates of control undamaged retinas, and retinas at different times after NMDA-treatment. Seurat was used to construct gene lists for differentially expressed genes (DEGs), violin/scatter plots, and dot plots. Significance of difference in violin/scatter plots was determined using a Wilcoxon Rank Sum test with Bonferroni correction. Genes that were used to identify different types of retinal cells included the following: (1) Müller glia: *GLUL, VIM, SCLC1A3, RLBP1*, (2) microglia: *C1QA, CSF1R, APOE*, *AIF1* (3) ganglion cells: *THY1, POU4F2, RBPMS2,* (4) amacrine cells: *GAD67, PAX6, TFAP2A*, (5) horizontal cells: *PROX1, CALB2,* (6) bipolar cells: *GRIK1, OTX1*, and (7) cone photoreceptors: *GNAT2, OPN1LW*, (8) rod photoreceptors: *RHO, NR2E3, ARR1,* (9) oligodendrocytes: *OLIG2, FGFR2, PMP2*, and (10) NIRG cells: *PTPRZ1, NKX2.2*. Activated MG were identified based on elevated expression levels of *NES, TGFB2, PMP2* and *MDK,* and MGPCs were identified based on expression of *CDK1, TOP2A, SPC25 and PCNA*, as described previously (Campbell et al., 2021c). The original detailed analysis of these scRNA-seq libraries are described elsewhere (Hoang et al., 2020; Campbell et al., 2021c; b, 2022).

For scATAC-seq, FASTQ files were processed using CellRanger ATAC 2.1.0, annotated using the chick genome (Gallus_gallus-5.0, Ensembl release 84, assembly GRC6a v105), and then aggregated using CellRanger ATAC aggr to create a common peak set. CellRanger ATAC aggr output was processed and analyzed using Signac (develop version 1.6.0.9015, github.com/timoast/signac) (Stuart et al., 2021) as described elsewhere (https://stuart.org/signac/). Annotations were added as an AnnotationHub (Bioconductor.org) object generated from the Ensembl chicken genome (see above). For Signac and chromVAR motif analysis, the genome used was a FaFile generated using Rsamtools (Bioconductor.org) and the necessary chicken FASTA and index files from Ensembl. For scATAC-seq analysis, including motif enrichment and motif scoring, all reference or annotation genomes were in NCBI/Ensembl format. The original detailed analysis of these scATAC-seq libraries are described elsewhere (Campbell et al., 2023).

### Photography, immunofluorescence measurements, and statistics

Wide-field photomicroscopy was performed using a Leica DM5000B microscope equipped with epifluorescence and Leica DC500 digital camera or Zeiss AxioImager M2 equipped with epifluorescence and Zeiss AxioCam 305 Mono. Confocal images were obtained using a Leica SP8 imaging system at the Department of Neuroscience Imaging Facility at the Ohio State University. Images were optimized for color, brightness and contrast, multiple channels over-laid, and figures constructed using Adobe Photoshop 2023. Cell counts were performed on representative images. To avoid the possibility of region-specific differences within the retina, cell counts were consistently made from the same region of retina for each data set.

Similar to previous reports (Fischer et al., 2009a; b; Ghai et al., 2009), immunofluorescence was quantified by using Image J (NIH). Identical illumination, microscope and camera settings were used to obtain images for quantification. Measurements of p21^Cip1^ and cFos immunofluorescence in the nuclei of MG were made for a fixed, cropped area of INL or INL+ONL. Measurements in the nuclei of MG were made by selecting the total area of pixels above threshold for Sox9 (in the red channel), copying p21^Cip1^ or cFos (in the green channel) into new images and measuring pixels intensities. The cropped areas contain between 50 and 150 MG or MGPCs; numbers of cells vary depending on treatments that influence the proliferation of MGPCs. Measurements were made for regions containing pixels with intensity values of 70 or greater (0 = black and 255 = saturated). The intensity sum was calculated as the total of pixel values for all pixels within threshold regions. GraphPad Prism 6 was used for statistical analyses.

A Levene’s test was used to determine whether data from control and treatment groups had equal variance. For treatment groups where the Levene’s test indicated unequal variance, a Mann Whitney U test (Wilcoxon Rank Sum Test) was used. For statistical evaluation of parametric data we used a two-tailed paired *t*-test to account for intra-individual variability where each biological sample served as its own control (left eye – control; right eye – treated). For multivariate analysis across >2 treatments, an ANOVA with the associated Tukey Test was performed to evaluate significant differences between multiple groups.

## Results

### Expression of ID transcription factors in embryonic retinas

We began by probing for patterns of expression of *ID1, ID2, ID3* and *ID4* in scRNA-seq libraries established from embryonic chick retinas at different stages of development, as described in previous reports (Campbell et al., 2021c, 2022; El-Hodiri et al., 2022). scRNA-seq libraries were established for retinal cells at embryonic day 5 (E5), E8, E12, and E15. These libraries included sequencing data for 22,698 cells after filtering to exclude doublets, cells with low UMI/genes per cell, and high mitochondrial gene-content. UMAP plots of aggregated libraries revealed clusters of cells that correlated to developmental stage and cell type (Fig. 1a). Types of cells were identified based on expression of well-established markers. Retinal progenitor cells (RPCs) from E5 and E8 retinas were identified by expression of *ASCL1*, C*DK1*, and *TOP2A*. (Fig. 1b). Maturing MG were identified by expression of *GLUL*, *RLBP1* and *SLC1A3* (Fig. 1b). *ID1*, *ID2* and *ID4* were expressed by maturing MG (mMG; Fig. 1c). In addition, *ID2* was detected in maturing bipolar cells (Fig. 1c). *ID3* was not widely expressed by cells in the embryonic chick retina (Fig. 1c). We bioinformatically isolated RPCs, immature MG (iMG) and mMG based on patterns of expression for *CDK1* and *TOP2A* (RPCs), *FABP7* and *VIM* (iMG), *GLUL,* and *RLBP1* (mMG) (Fig. 1d-g). We found that *ID1* and *ID4* were expressed at significantly elevated levels by mMG from E12 and E15 retinas (Fig. 1h). By comparison, *ID2* upregulated iMG and mMG at E18 and E12 in the developing retina, whereas *ID3* was not widely expressed by RPCs, iMG and mMG (Fig. 1h).

**Figure 1:**
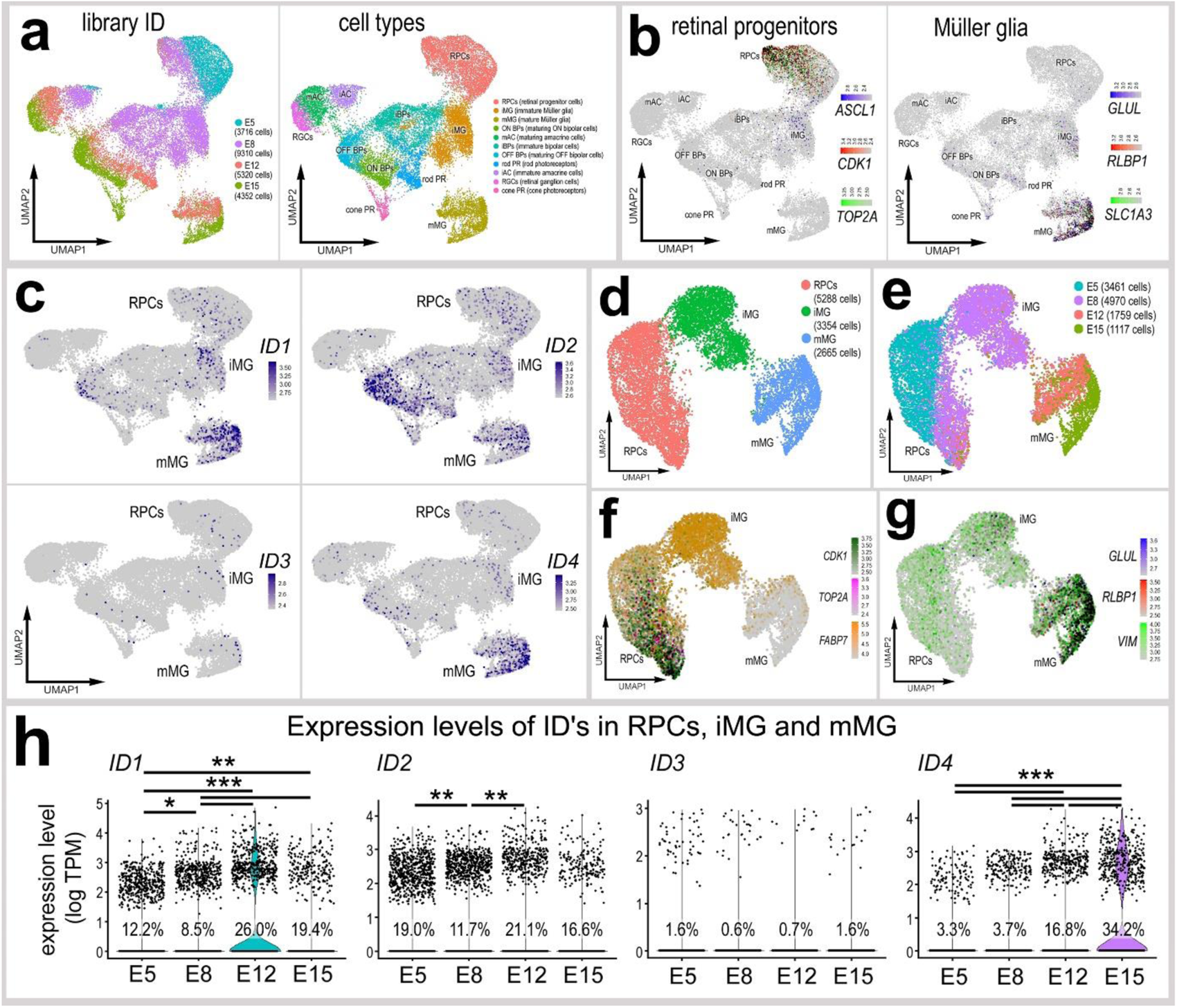
Patterns of expression of ID TFs in embryonic retina: scRNA-seq libraries were generated from embryonic retinal cells at E5, E8, E12 and E15 (**a**). UMAP plots indicate cell types that were identified by expression of cell-distinguishing genes for progenitors or mature glia, with cells expressing 2+ genes denoted in black (**b**). Expression patterns of *ID1, ID2, ID3* and *ID4* are illustrated in UMAP heatmaps (**c**). RPCs, iMG and mMG were bioinformatically isolated and re-embedded in UMAP plots (**d-g**). Violin plots illustrate expression levels of ID isoforms in UMAP-clustered RPCs and MG (**h**). (**h**) *p<0.01, **p<0.001, ***p<<0.0001; Wilcoxon Rank Sum Test with Bonferroni correction. RPC – retinal progenitor cell, MG – Müller glia, iMG – immature Müller glia, mMG – mature Müller glia.

### Expression of ID transcription factors in damaged or FGF-treated mature retinas

We next probed for patterns of expression of *ID1, ID2, ID3* and *ID4* in scRNA-seq libraries established for post-hatch chick retinas, as described in previous reports (Campbell et al., 2019; Todd et al., 2019; Campbell et al., 2021b, 2022). scRNA-seq libraries were established for cells from control retinas and retinas at 24, 48 and 72hrs after NMDA-treatment (Fig. 2a). Retinal cell types ordered into UMAP clusters were identified based on well-established markers (Fig. 2b), as described in the methods. MGPCs were identified based on patterns of expression of *SPC25, PCNA, TOP2A* and *CDK1* (Fig. 2c). We found that *ID1* was expressed by relatively few amacrine and bipolar cells, and by MGPCs and MG returning to a resting phenotype (Fig. 2d). *ID2* was expressed by bipolar and amacrine cells, cone photoreceptors and NIRG cells, as well as a few MGPCs (Fig. 2d). *ID3* was expressed by relatively few scattered cells including bipolar cells and activated MG (Fig. 2d). *ID4* was highly expressed by resting MG, appeared to be downregulated in activated MG at 24hrs after NMDA and in MGPCs, and appeared to be upregulated in MG returning to resting (Fig. 2d).

**Figure 2:**
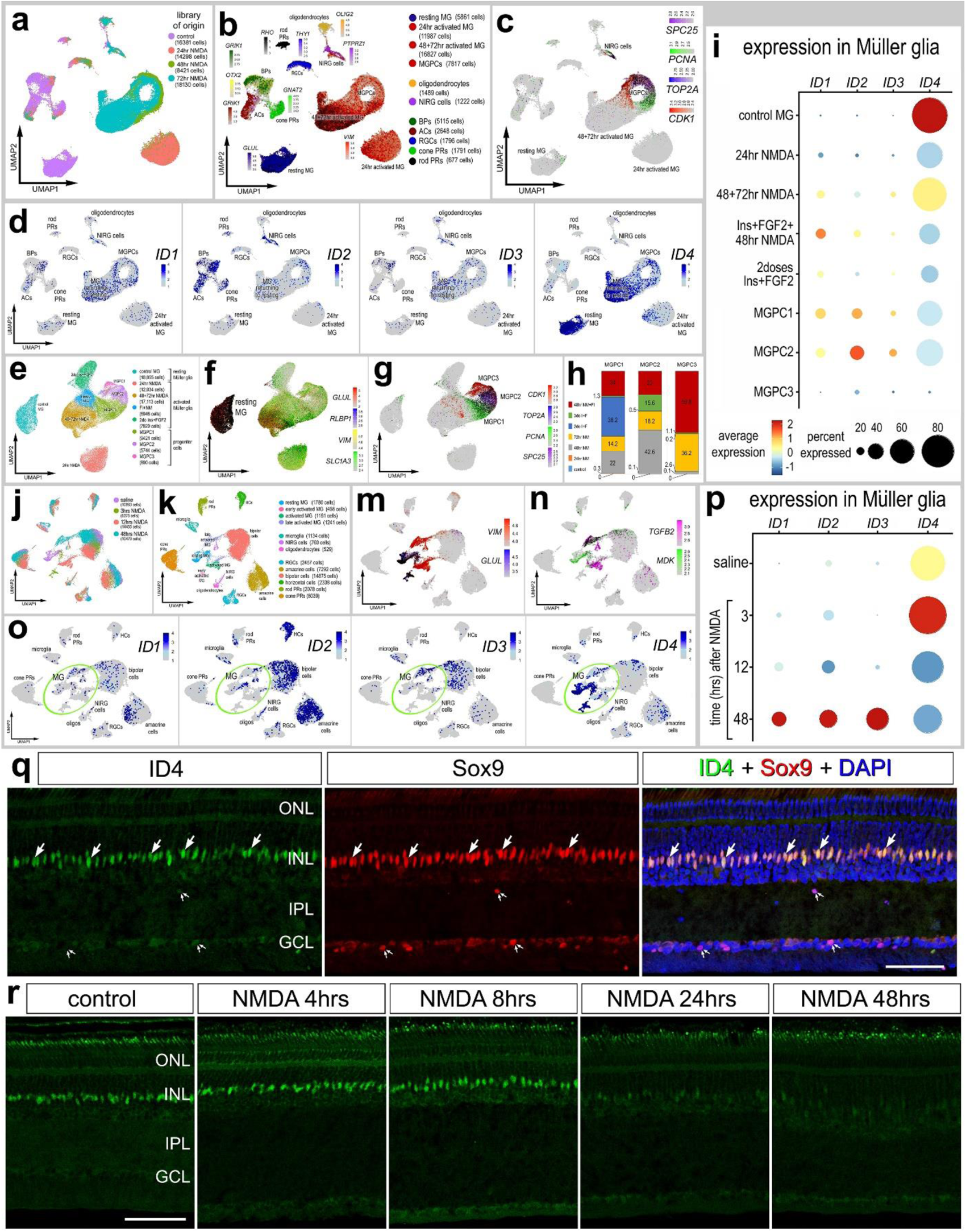
Patterns of expression of ID TFs in mature retinas. scRNA-seq was used to identify patterns of expression of ID TFs among retinal cells with the data presented in UMAP (**a**-**g, j-o**) or dot plots (**i,p**). Aggregate scRNA-seq libraries were generated for cells from; (i) control retinas and retinas 24, 48, and 72hrs after NMDA-treatment (**a-d**), (ii) MG were bioinformatically isolated from control retinas, retinas at 24, 48 and 72hrs times after NMDA, retinas treated with 2 or 3 doses of insulin and FGF2, and retinas treated with insulin, FGF2 and NMDA (**e-i**), and (iii) control retinas and retinas 3, 12 and 48hrs after NMDA (**j-p**). UMAP-ordered cells formed distinct clusters of neuronal cells, resting MG, early activated MG, activated MG and MGPCs. Resting MG were identified based on elevated expression of *SLC1A3, RLBP1, VIM* and *GLUL* (**b,f**), activated MG down-regulated *SLC1A3, RLBP1* and *GLUL*, and MGPCs up-regulated proliferation markers including *SPC25, PCNA, TOP2A* and *CDK1* (**c,g**). UMAP heatmap plots for *ID1, ID2, ID3* and *ID4* illustrate levels and patterns of expression across all retinal cell types (**d, o**). MGPCs were comprised of cells from different treatment groups; predominantly cells from retinas at 72 h after NMDA-treatment and 48 hrs after NMDA+insulin+FGF2 (**g,h**). Dot plots illustrate relative levels of expression and percent expression of *ID1, ID2, ID3* and *ID4* in MG in different UMAP clusters (**i**) or library of origin (**p**). Immunolabeling for ID4 in normal and NMDA-damaged retinas at 4, 8, 24 and 48hrs after treatment, with treatment starting at P10 (**q,r**). Images were taken from mid-peripheral regions of the retina. Arrows indicate the nuclei of MG. The calibration bars in **q** and **r** represents 50 µm. Abbreviations: ONL – outer nuclear layer, INL – inner nuclear layer, IPL – inner plexiform layer, GCL – ganglion cell layer.

To gain a more comprehensive understanding of patterns of expression of ID TFs, we probed a large aggregate scRNA-seq library of MG (>70,000 MG and MGPCs) with glia bioinformatically isolated from retinas treated with saline, 24/48/72hrs NMDA, 2 or 3 doses of insulin+FGF2 and 48hrs NMDA+insulin+FGF2 (Fig. 3e), as described previously (Campbell et al., 2021c, 2022; El-Hodiri et al., 2022). Resting MG and activated MG at 24hrs after NMDA formed 2 discrete clusters of cells, whereas activated MG from different treatment groups and MGPCs formed a continuum of cells based, in part, on expression of cell cycle regulators (Fig. 2f-h). In resting MG, relative levels of *ID4* were very high whereas levels of *ID1, ID2* and *ID3* were very low (Fig. 2i). After treatment with NMDA and/or insulin+FGF2 levels of *ID4* were significantly reduced, with exception to elevated levels of *ID4* in MG at 48 and 72 hrs after NMDA from the cluster of MG that were returning toward a resting phenotype (Fig. 2i). Levels of *ID1, ID2* and *ID3* were highest in MGPC1 and MGPC2 clusters, whereas levels of all ID factors were very low in the MGPC3 cluster and in the MG treated with 2 doses of insulin+FGF2 (Fig. 2i).

**Figure 3:**
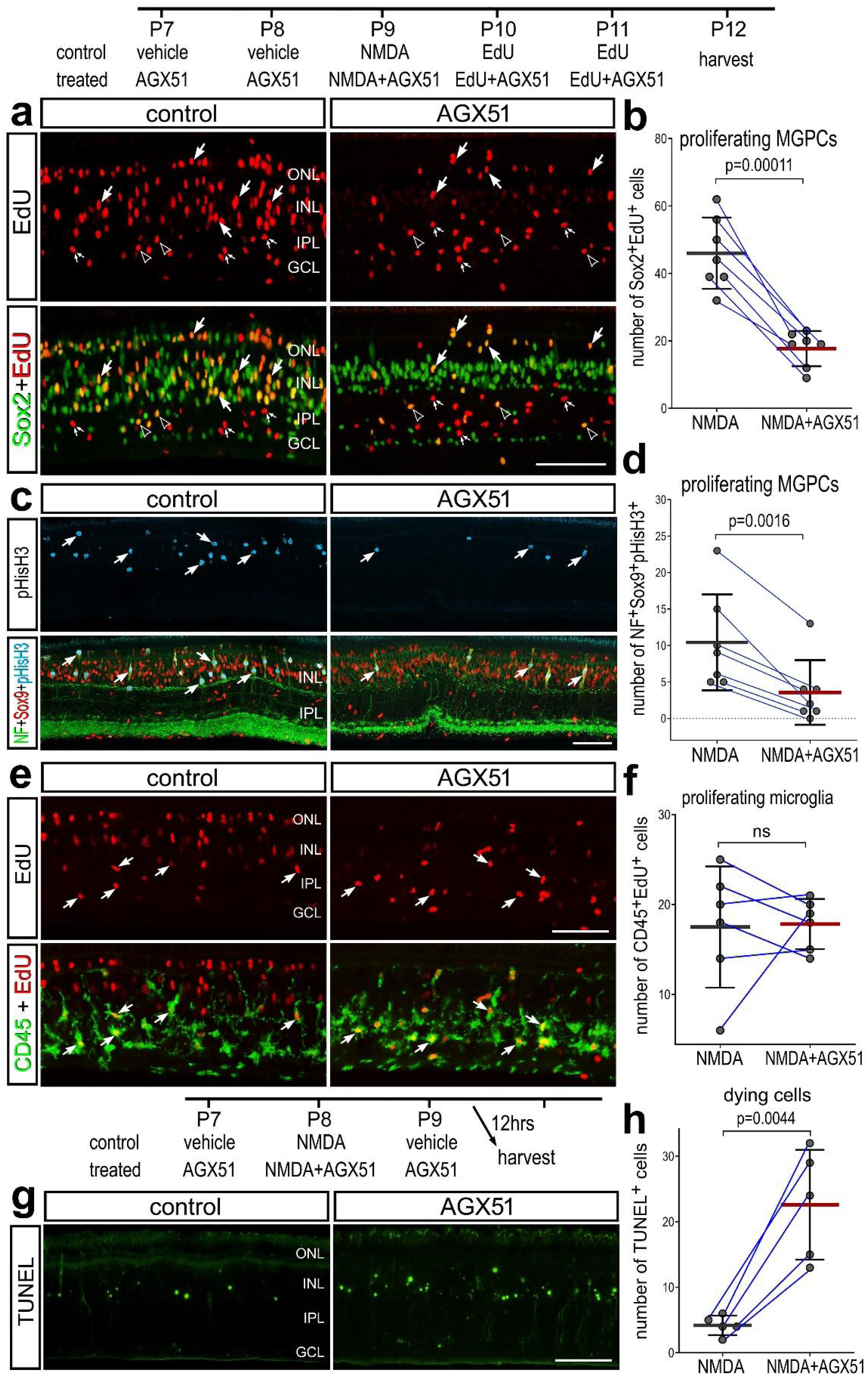
Inhibition of ID TFs suppresses the proliferation of MGPCs. Chick eyes were injected with vehicle or AGX51 at P7 and P8, NMDA ± AGX51 at P9, EdU ± AGX51 at P10 and harvested at P11. Retinas sections were labeled for EdU (red; **a,e**), antibodies to Sox2 (green; **a**), phospho-histone H3 (blue; **c**), neurofilament (green; **c**), Sox9 (red; **c**), CD45 (green; **e**) or fragmented DNA (TUNEL, green; **g**). Histograms (**b,d,f,h**) represent the mean (bar ± SD) and each dot represents one biological replicate (blue lines connect data points from control and treated retinas from the same individual). Significance of difference (p-values) was determined by using a paired t-test. Calibration bars in **a**,**c**,**e** and **g** represent 50 µm. Arrows indicate the nuclei of MGPCs (**a,c)** or the nuclei of microglia (**e**). Abbreviations: ONL – outer nuclear layer, INL – inner nuclear layer, IPL – inner plexiform layer, GCL – ganglion cell layer, ns – not significant.

We further analyzed expression of ID TFs in a scRNA-seq database that we generated for early time-points, 3 and 12 hours, after NMDA-treatment (Campbell et al., 2021b, 2023) (Fig. 2j). These scRNA-seq libraries were generated with different reagents and sensitivities. Thus, they were not integrated with the older scRNA-seq libraries, and were analyzed separately. UMAP ordering of cells revealed distinct clusters of neurons and glia, with MG forming distinct clusters based on time after NMDA-treatment (Fig. 2j-n). Patterns of expression for ID factors among retinal neurons and glia were consistent between these different libraries, with the exception of expression in microglia and horizontal cells which were well represented in libraries from early time point after NMDA (Fig. 2o). We found that relative levels of *ID4* were significantly increased in MG at 3hrs after NMDA-treatment, and then rapidly decreased by 12 hrs after NMDA (Fig. 2o-p). By comparison, levels of *ID1, ID2* and *ID3* were elevated only in MG from retinas at 48hrs after NMDA-treatment (Fig. 2o-p).

To validate findings from scRNA-seq, we applied antibodies to ID4 to sections of retinas collected from post-hatch day 10 (P10) chicks. In undamaged retinas, ID4 immunoreactivity was detected in the nuclei of Sox9-positive MG (Fig. 2q). In damaged retinas, levels of ID4 immunoreactivity remained high at 4 and 8hrs after NMDA-treatment (Fig. 2r). Levels of ID4 were greatly reduced at 24hrs post-injury and were nearly absent at 48hrs (Fig. 2r). These patterns of immunolabeling are consistent with patterns of expression of *ID4* seen in scRNA-seq libraries. We applied different antibodies to ID1, ID2 and ID3, but none of these antibodies produced plausible patterns of labeling (not shown).

### Inhibition of ID transcription factors in damaged retinas

Since ID4 was predominantly expressed by resting MG in undamaged retinas, we applied an Id inhibitor (AGX51) for 2 consecutive days prior to treatment with NMDA, with NMDA, and 2 consecutive days following NMDA-treatment. AGX-51 interferes with ID interactions with target bHLH protein E47(*TCF3*; (Wojnarowicz et al., 2021). Further, AGX51 has been shown to drive ubiquitin-mediated degradation of ID proteins. We found that the ID inhibitor significantly decreased numbers of MGPCs labeled for Sox2 and EdU in NMDA-damaged retinas (Fig. 3a,b). Consistent with these findings, we found signficant decreases in numbers of MGPCs labeled for neurofilament, Sox9 and pHisH3 in damaged retinas treated with AGX-51 (Fig. 3c,d). However, the ID inhibitor had no significant effect upon numbers of proliferating microglia (Fig. 3e,f). Further, we found that treatment with AGX-51 significantly increased numbers of dying cells (Fig. 3g,h). Collectively, these findings indicate that ID factors suppress the proliferation of MGPCs in damaged retinas and support neuronal survival.

### Inhibition of ID transcription factors and neurogenesis

We next investigated whether ID TFs regulate the neurogenic potential of MGPCs in the chick retina. During embryonic development, IDs are known to promote glial cell fate (reviewed by (Ross et al., 2003). In mature, damaged mouse retinas inhibition of IDs has been shown to increase the differentiation of bipolar-like cells from Ascl1-over expressing MG (Palazzo et al., 2022). Further, we find that levels of *ID1, ID2* and *ID3* were elevated in MGPCs in the chick retina (see Fig. 2), consistent with the notion that these factors suppress the neurogenic potential of MGPCs. Accordingly, we tested whether inhibition of IDs, starting at 2 days after NMDA-treatment when MG have committed to re-enter the cell cycle, influences the differentiation of progeny. We found that AGX51 had no significant effects upon proliferation of MGPCs when applied 2, 3, and 4 days NMDA (Fig. 4a,b,c). We found a significant increase in numbers of EdU-labeled amacrine-like cells that were co-labeled for HuC/D (Fig. 4d,e) or calretinin (Fig. 4f,g). In addition, we found a small, but significant, increase in numbers of EdU-labeled amacrine-like cells that were co-labeled for Ap2a (Fig. 4h). These results indicate that ID activity following NMDA damage normally suppresses the neurogenic potential of MGPCs.

**Figure 4:**
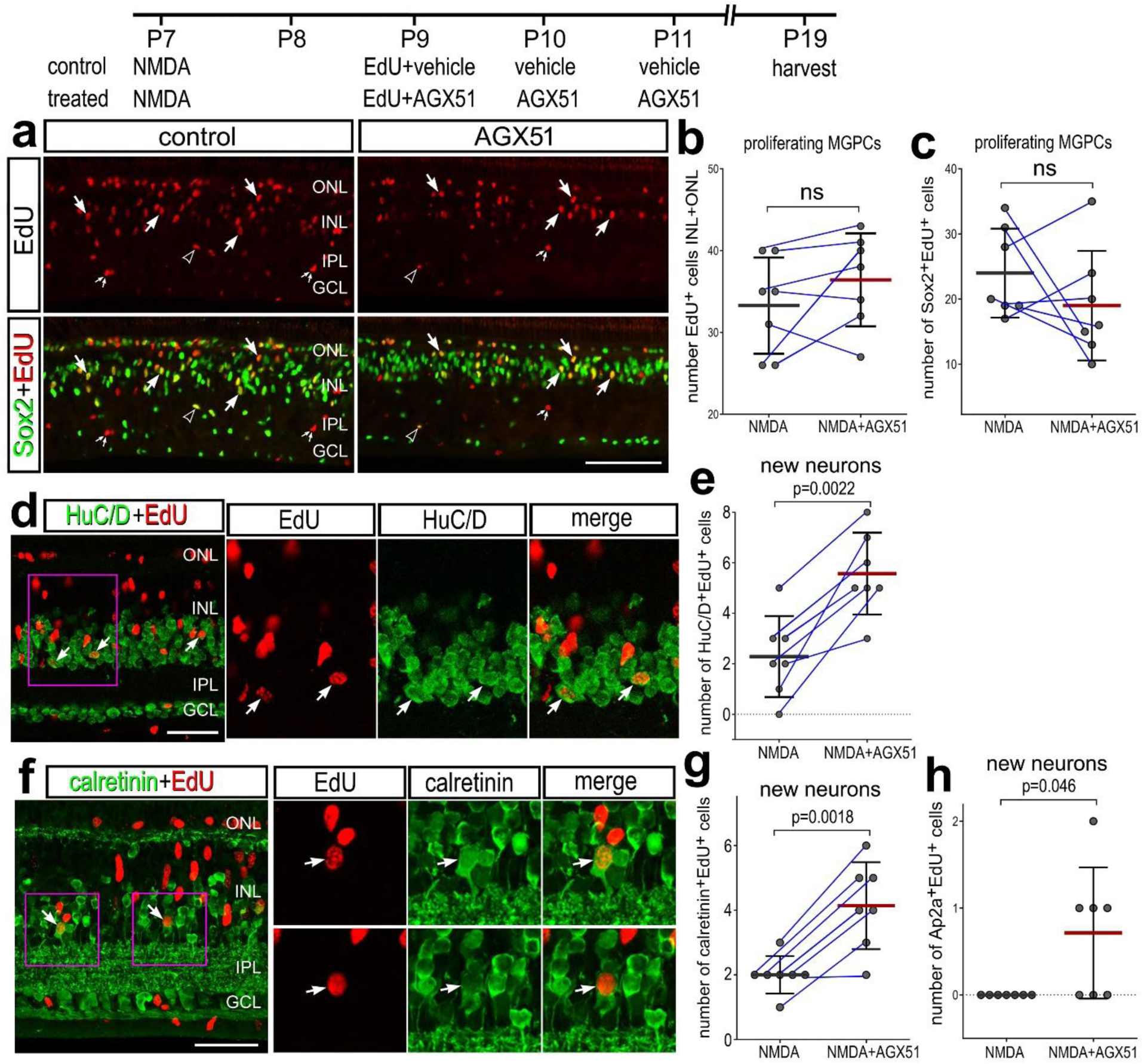
Inhibition of ID TFs increases neuronal differentiation of MGPC progeny. Chick eyes were injected with NMDA at P7, vehicle ± AGX51 at P9, P10 + P11, and harvested at P12. Retinas sections were labeled for EdU (red; **a,d,g**) and antibodies to Sox2 (green; **a**), HuC/D (green; **d**), or calretinin (green; **f**). Histograms represent the mean (bar ± SD) and each dot represents one biological replicate (blue lines connect control and treated eyes from the same individual) for proliferating MGPCs (**b,c**) or newly generated neurons (**e,g,h**). Significance of difference (p-values) was determined by using a paired t-test. Regions of interest (magenta boxes) are enlarged 1.8-fold and displayed in separate channels. Arrows indicate nuclei of MGPCs (**a**) or newly generated neurons (**d,f**), small double-arrows indicate the nuclei of putative microglia, and hollow arrowheads indicate the nuclei of NIRG cells. Calibration bars represent 50 µm (**a**,**d**,**f**). Abbreviations: ONL – outer nuclear layer, INL – inner nuclear layer, IPL – inner plexiform layer, GCL – ganglion cell layer, ns – not significant.

### Inhibition of ID transcription factors in undamaged retinas

In the chick retina, the formation of MGPCs can be stimulated in the absence of neuronal damage in response to consecutive daily injects of insulin and FGF2 (Fischer et al., 2002) or FGF2 alone (Fischer et al., 2014b). Accordingly, we sought to test whether ID inhibition influenced the formation of MGPCs in undamaged retinas treated with 3 consecutive doses of insulin and FGF2. We found no significant difference in numbers of proliferating MGPCs that were labeled for Sox2 and EdU in response to AGX51 pretreatment (Fig. 5a,b). We detected very few dying cells in retinas treated with insulin+FGF2, and ID inhibitor did not influence numbers of dying cells (Fig. 5c). Similarly, we observed small but significant increases in numbers of proliferating microglia in retinas treated with insulin, FGF2 and ID inhibitor, without affecting levels of CD45 (Fig. 5d,e,f).

**Figure 5:**
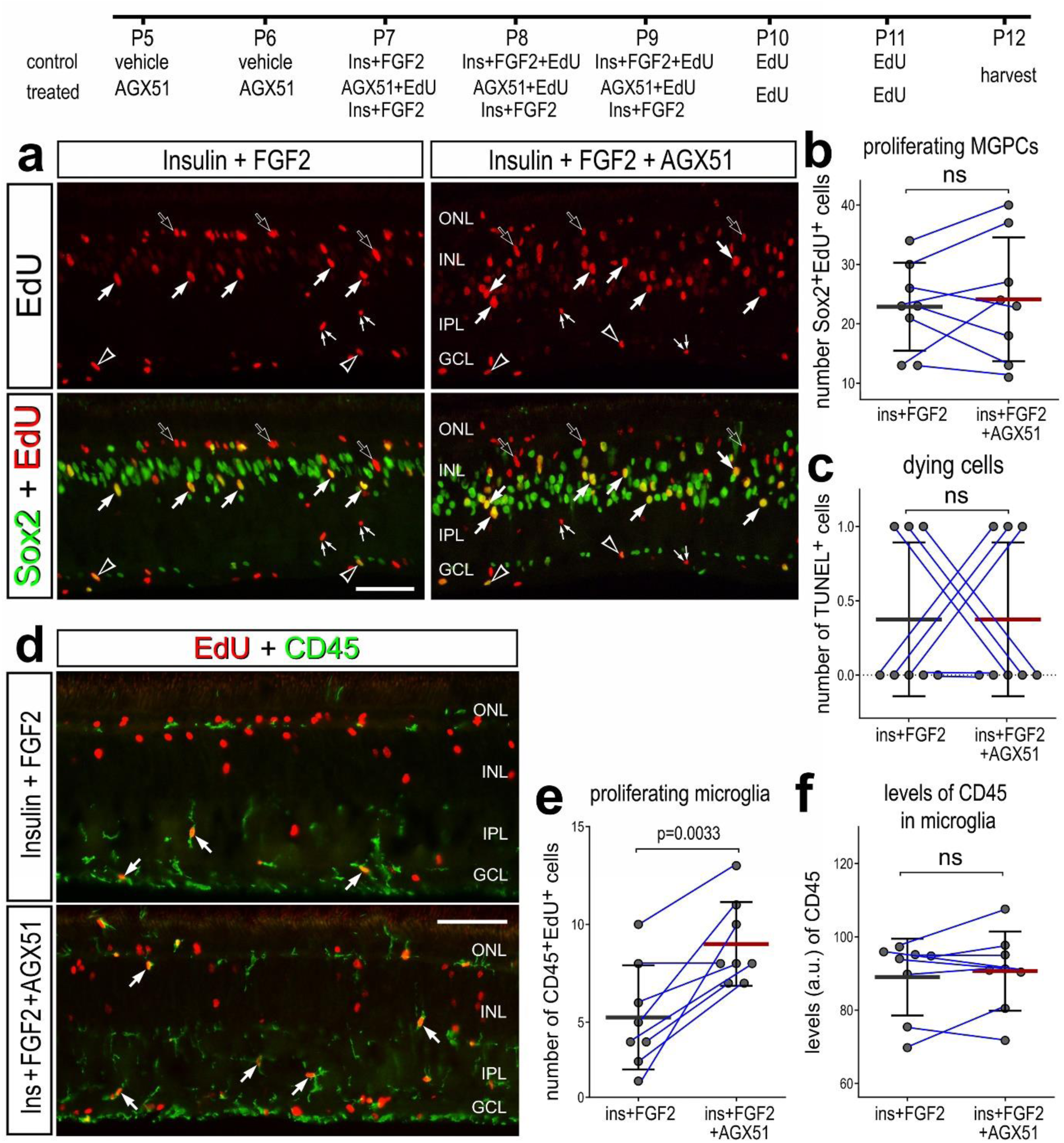
Inhibition of ID TFs does not affect the formation of MGPCs in undamaged retinas. Chick eyes were injected with saline ± AGX51 at P5 and P6, insulin and FGF2 ± AGX51 at P7, P8 and P9, EdU at P10 and P11, and retinas harvested at P12. Retinal sections were labeled for EdU (red; **a**) and antibodies to Sox2 (green; **a**), or CD45 (green; **d**). Histograms represent the mean (bar ± SD) and each dot represents one biological replicate (blue lines connect data points from control and treated retinas from the same individual) for proliferating MGPCs labeled for Sox2 and EdU (**b**) dying cells that were TUNEL-positive (**c**), proliferating microglia labeled for CD45 and EdU (**e**), and CD45 levels (**f**) Significance of difference (p-values) was determined by using a paired t-test. The calibration bars in **a** and **d** represent 50 µm. Abbreviations: ONL – outer nuclear layer, INL – inner nuclear layer, IPL – inner plexiform layer, GCL – ganglion cell layer, ns – not significant.

To better understand why ID inhibition had no effect on the formation of proliferating MGPCs in retinas treated with insulin+FGF2, we probed scRNA-seq libraries for levels and patterns of expression of ID factors. Treatment with insulin and FGF2 potently impacts patterns of gene expression in MG (Hoang et al., 2020; Campbell et al., 2021b, 2022). UMAP ordering of cells provided distinct clusters of retinal neurons from all treatment groups, whereas MG from different treatment groups formed distinct clusters of cells (Fig. 6a-c), indicating treatment with 2 or 3 doses of insulin+FGF2 broadly impacted gene expression in MG, but not neurons. For example, genes expressed by resting MG, including *GLUL* and *RLBP1*, are robustly downregulated and some genes expressed by activated MG, including *PMP2* and *S100A6*, are robustly upregulated in response to insulin and FGF2 (Fig. 6d,e). Further MGPCs were found only in retinas treated with 3 doses of insulin+FGF2 and formed a sub-cluster in UMAP plots, identified by upregulation of *CDK1* and *TOP2A* (Fig. 6c,f). There was a modest, but significant (p<0.0001), upregulation of *ID1* and *ID2* in MG and MGPCs in retinas treated with 3 doses of insulin and FGF2 (Fig. 6g,h,k). *ID3* was not widely expressed by neurons and glia in retinas treated with saline or insulin and FGF2 (Fig. 6i,k). By comparison, the high levels of *ID4* in resting MG were dramatically downregulated in MG from retinas treated with 2 or 3 doses of insulin and FGF2 (Fig. 6j,k). To validate these findings we applied antibodies to ID4 to retinas treated with different doses of insulin+FGF2. Consistent with data from scRNA-seq analyses, 3 doses of insulin+FGF2 greatly diminished ID4 immunofluorescence (Fig. 6l). Similarly, we found that a single dose of insulin+FGF2 significantly reduced levels of ID4 in the MG (Fig. 6m). These findings suggest that AGX51 may have had no effect on MGPC formation because Insulin and FGF rapidly downregulate ID4 in MG, unlike NMDA-induced damage which rapidly upregulated levels of *ID4*.

**Figure 6:**
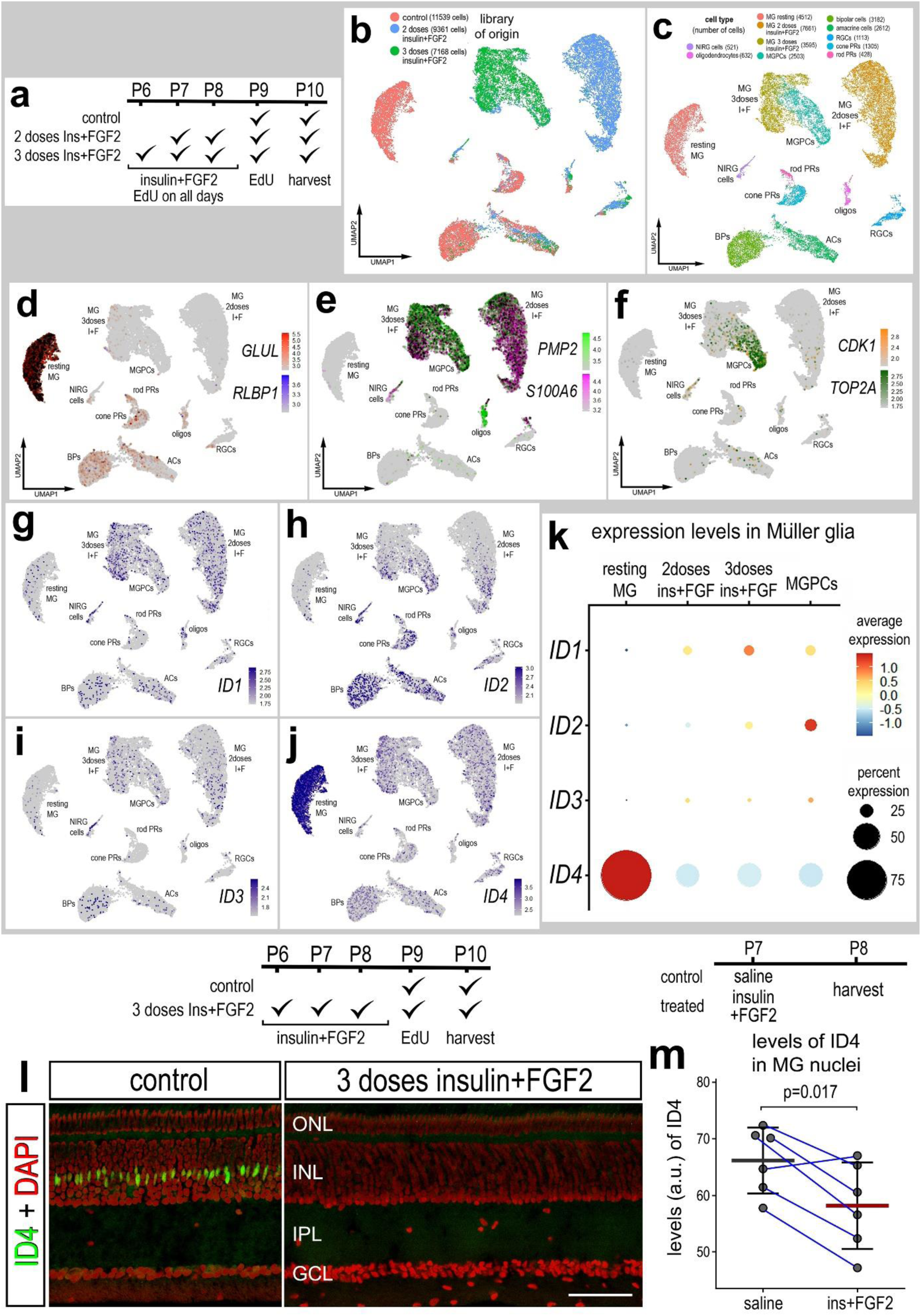
Treatment with insulin and FGF24 downregulates levels of ID TFs. We probed for expression patterns of ID TFs in scRNA-seq libraries from retinas treated vehicle, or with 2 or 3 doses of insulin and FGF2 (**a**). Libraries contained between 7100 and 11600 cells wherein different types of neurons form distinct clusters regardless of treatment, and MG formed different distinct clusters based on treatment (**b**,**c**). Resting MG from control retinas were identified based on high levels of expression of *GLUL* and *RLBP1* (**d**), activated MG identified based on expression of PMP2 and S100A6 (**e**), and MGPCs identified based on expression of CDK1 and TOP2A (**f**). UMAP heatmap plots illustrate patterns and levels of expression of *ID1, ID2, ID3* and *ID4* in MG (**g,h,i,j**). The dot plot in **k** illustrates levels and percent expression of *ID1, ID2, ID3* and *ID4* in MG. Significance of difference (***p<<0.0001) was determined using a Wilcoxon Rank Sum Test with Bonferroni correction. (**l**) retinal sections were label with DAPI (red nuclei) and antibodies to ID4 (green). The calibration bar in **l** represents 50 µm. Abbreviations: ONL – outer nuclear layer, INL – inner nuclear layer, IPL – inner plexiform layer, GCL – ganglion cell layer, ns – not significant. The histogram in **m** represents the mean (bar ± SD) and each dot represents one biological replicate (blue lines connect data points from control and treated retinas from the same individual) for levels of ID4 in the nuclei of MG.

### ID transcription factors regulate cyclin dependent kinase inhibitor expression

ID TFs are known to have many different transcriptional targets, including cyclin dependent kinase inhibitors, including p21^Cip1^ (Carey et al., 2009; Sharma et al., 2012; Knowell et al., 2013). Thus, we characterized patterns of expression of *CDKN1A* (p21^Cip1^) in different scRNA-seq libraries. In scRNA-seq libraries of embryonic retinas, we observed that *CDKN1A* was upregulated by maturing MG from E12 and E15 retinas, but was not widely expressed by other types of cell in embryonic chick retinas (Fig. 7a, see Fig). This pattern of expression was very similar to that seen for *ID4* in developing retinas (Fig. 1a-b for legend). In post-hatch chick retinas, *CDKN1A* was expressed at high levels by resting MG and was significantly downregulated in activated MG at 24hrs after NMDA and in MG (Fig. 7b; see Fig. 2a-c for legend). Levels of *CDKN1A* were elevated in MG returning to a resting state from retinas at 48 and 72 hours after NMDA-treatment (Fig. 7b). In the absence of retinal damage, *CDKN1A* was greatly reduced by 2 doses of insulin and FGF2, further reduced by 3 doses of insulin and FGF2 and nearly absent in MGPCs (Fig. 7c, see Fig 6a-c for legend). In an aggregate database of MG from post-hatch retinas from different treatments, the largest downregulation of *CDKN1A* was observed at 24hrs after NMDA and in MGPCs from FGF+Insulin or NMDA treatment (Fig. 7d; see Fig. 2e-h for legend).

**Figure 7:**
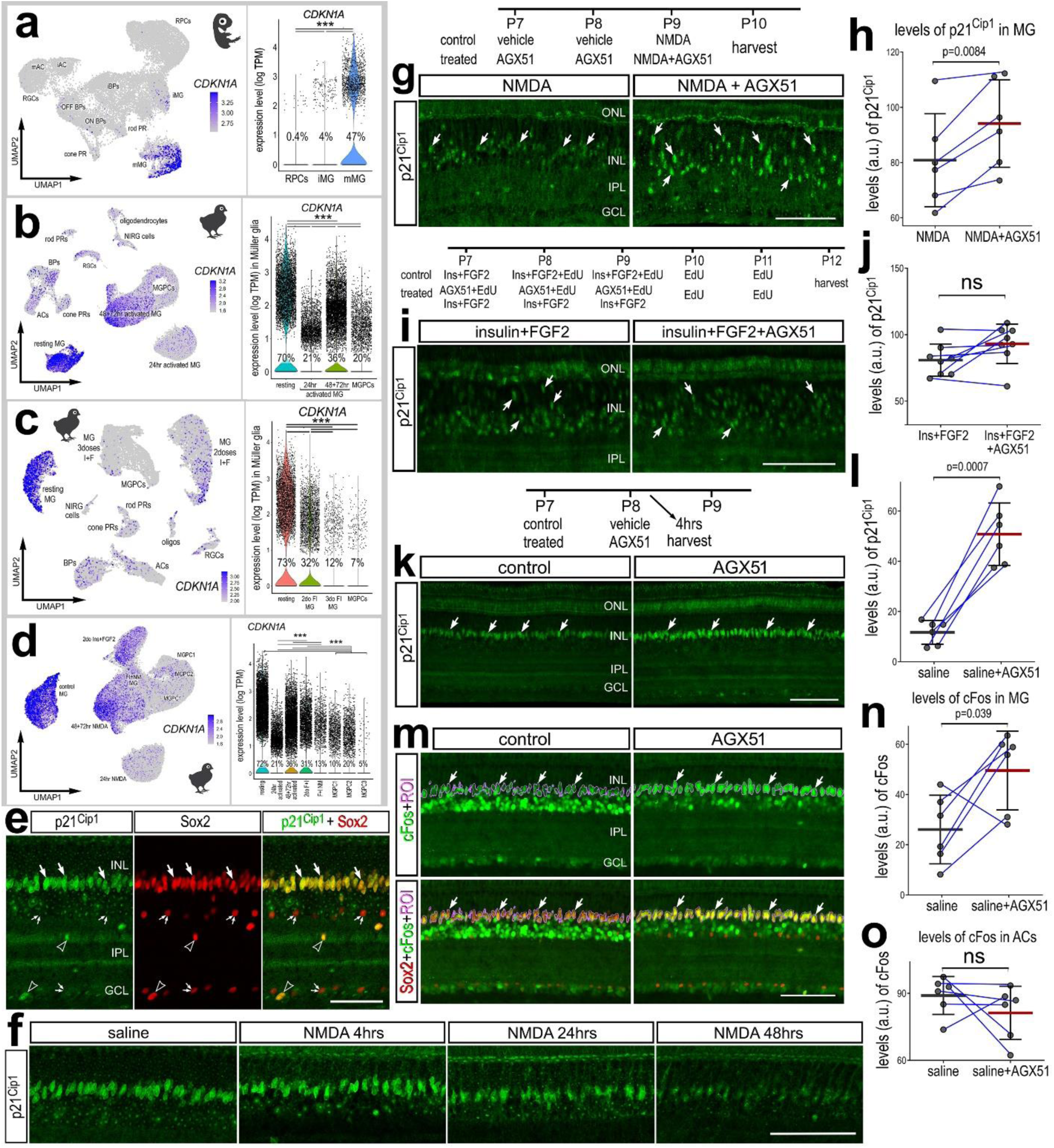
Patterns of p21^Cip1^ (*CDKN1A*) expression in maturing, resting MG and activated MG. We probed for patterns of expression scRNA-seq libraries from embryonic retinas (**a**), retinas treated with insulin and FGF2 (**b**) NMDA-damaged post-hatch retinas (**c**), and aggregated MG from retinas damaged by NMDA and/or treated with insulin and FGF2 (**d**). UMAP heatmap plots illustrate patterns and levels of expression of *CDKN1A.* Violin plots illustrate levels and percent expression of *CDKN1A* in MG. Significance of difference (*p<0.001, ***p<<0.0001) was determined using a Wilcoxon Rank Sum Test with Bonferroni correction. **Inhibition of ID TFs stimulate MG to upregulate p21^Cip1^ and cFos.** Retinas were obtained from untreated eyes (**e,f**) and eyes injected with NMDA (**f**,**g,h**), insulin and FGF2 ± AGX51 (**i,j**), or AGX51 alone (**i-m**). Sections of the retina were labeled with antibodies to p21^Cip1^ (green; **e,f,g,i**), Sox2 (red; **e,g,i,k**), or cFos (green; **m**). The magenta regions of interest in **m** outline the nuclei of Sox2+ MG. Arrows indicate double-labeled nuclei of MG, hollow arrowheads indicate the nuclei of NIRG cells and small double-arrows nuclei of cholinergic amacrine cells. The calibration bars represent 50 µm. Abbreviations: ONL – outer nuclear layer, INL – inner nuclear layer, IPL – inner plexiform layer, GCL – ganglion cell layer, ns – not significant. Histograms represent the mean (bar ± SD) and each dot represents one biological replicate (blue lines connect data points from control and treated retinas from the same individual) for levels of p21^Cip1^ (**h,j,l**) or cFos (**n,o**). P-values were determined by using a paired t-test.

To validate data from scRNA-seq, we applied antibodies to p21^Cip1^ to retinal sections from control and NMDA-treated eyes. We found p21^Cip1^ immunoreactivity in Sox2-positive MG nuclei in the middle of the INL (Fig. 7e). Further, we observed p21^Cip1^ immunoreactivity in Sox2-positive nuclei of NIRG cells scattered in the IPL and GCL, but not in the Sox2-positive nuclei of cholinergic amacrine cells in the INL and GCL (Fig. 7e). NIRG cells and cholinergic amacrine cells are known to express Sox2 (Fischer et al., 2010). Levels of p21^Cip1^ immunoreactivity in MG nuclei appeared high at 4hrs after NMDA-treatment, slightly reduced at 24hrs and nearly absent at 48hrs after treatment (Fig. 7f).

We next tested whether ID inhibition influenced levels on p21^Cip1^ and other factors in undamaged retinas. We found that treatment of NMDA-damaged retinas with ID inhibitor significantly increased levels of p21^Cip1^ in the nuclei of MG compared to levels seen in retinas treated with NMDA alone (Fig. 7g,h). We found that treatment of retinas with insulin, FGF2 and ID inhibitor had no significant effect on levels of p21^Cip1^ in the nuclei of MG compare to levels seen in retinas treated with insulin+FGF2 alone (Fig. 7i,j). In undamaged retinas, we found that a single intraocular injection of ID inhibitor significantly increase in levels of p21^Cip1^ in the nuclei of MG (Fig. 7k,l). Furthermore, we found that treatment with ID inhibitor significantly increased levels of immediate early gene cFos in the nuclei of MG, whereas levels of cFos in the nuclei of amacrine cells was not significantly affected (Fig. 7m-o). By comparison, treatment with AGX51 had no effect upon levels of pAkt or glutamine synthetase in MG (not shown).

### Notch- and gp130-signaling influence ID4 and p21Cip1

Notch signaling has been shown to regulate id2a expression in damaged zebrafish retinas (Luo et al., 2012). In the chick retina, Notch-signaling is known to be required for the proliferation of MGPCs in retina treated with NMDA or insulin+FGF2, and suppresses the neuronal differentiation of progeny (Hayes et al., 2007; Ghai et al., 2010). Similarly, signaling through gp130/Jak/Stat is required for the proliferation of MGPCs and suppresses the neuronal differentiation of progeny in the chick retina (Todd et al., 2016a). Accordingly, we tested whether inhibition of Notch or gp130 influenced the expression of ID4 or p21^Cip1^ in MG in damaged retinas. We found that inhibition of Notch with DAPT or inhibition of gp130 with sc144 in NMDA-damaged retinas resulted in significant increases in ID4 immunoreactivity in the nuclei of MG (Fig. 8a,b,c). Unlike our findings from ID inhibition, levels of p21^Cip1^ were not significantly affected by Notch or gp130 inhibitors (Fig. 8d,e,f). These data indicate that Notch and gp130/Jak/Stat signaling suppress ID4 expression in MG in damaged retinas.

**Figure 8:**
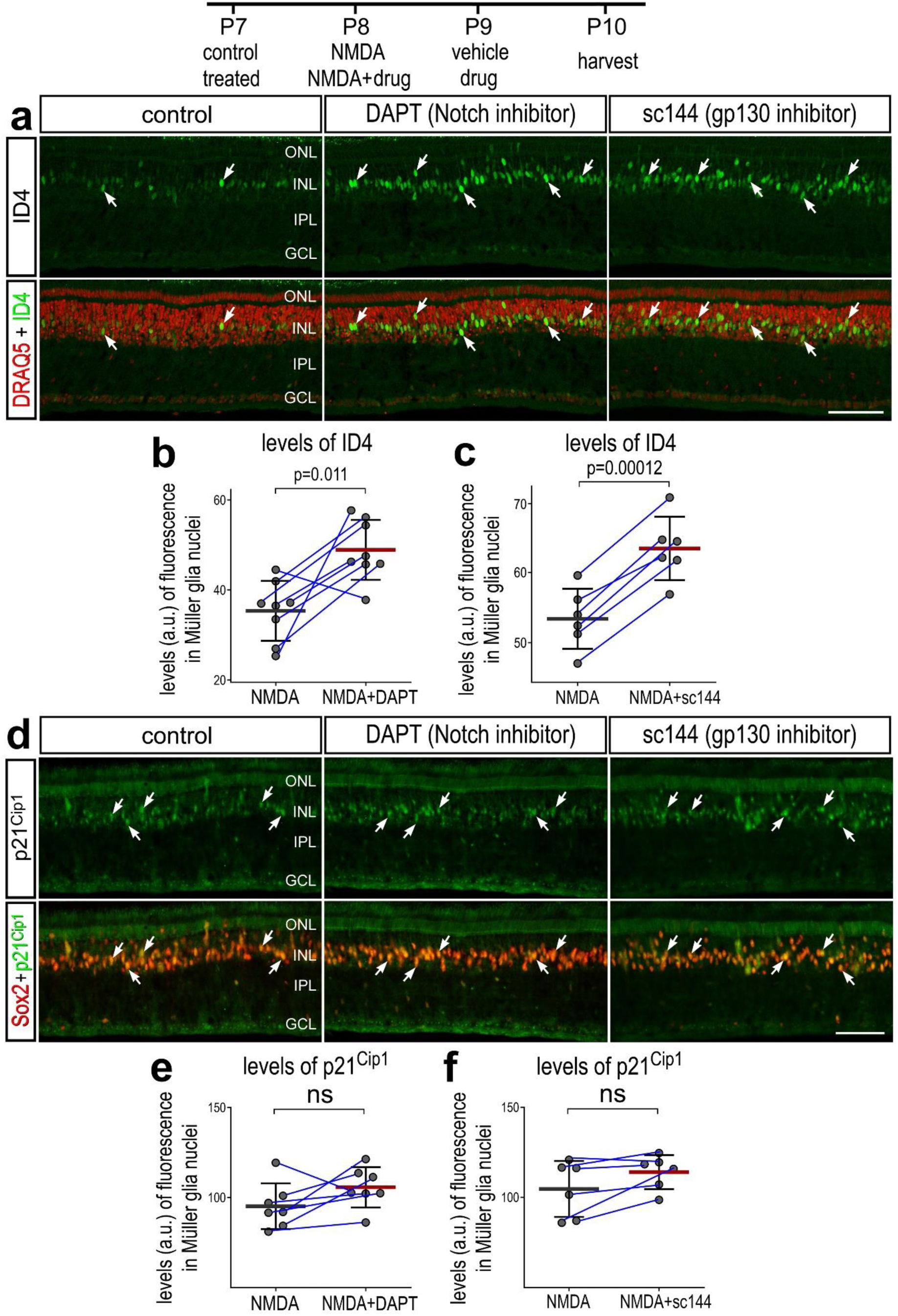
Notch- and gp130-signaling suppress ID4, but not p21^Cip1^ in MG in damaged retinas. Retinas were obtained from eyes injected with NMDA ± DAPT or ± sc144 at P8, vehicle or DAPT or sc144 at P9, and tissues harvested at P10. Sections of the retina were labeled with Draq5 (red; **a**) and antibodies to ID4 (green; **a**) or Sox2 (red; **d**) and p21Cip1 (green; **d**). Histograms (**b,c,e,f**) represent the mean (bar ± SD) and each dot represents one biological replicate (blue lines connect data points from control and treated retinas from the same individual). Significance of difference (p-values) was determined by using a paired t-test. Arrows indicate the nuclei of MG. Abbreviations: ONL – outer nuclear layer, INL – inner nuclear layer, IPL – inner plexiform layer, GCL – ganglion cell layer, ns – not significant. Calibration bars represent 50 µm.

### Changes in chromatin access for IDs and CDKN1A

We treated with saline or NMDA and harvested retinal cells 24hrs after injection to generate scATAC-seq libraries, as described previously (Campbell et al., 2023). UMAP embedding of cells revealed distinct clusters of different types of neurons and glia (Fig. 9a,b). We identified clusters of cell types from elevated chromatin access for genes known to be expressed by MG (*SOX2, NOTCH1, PAX6*), oligodendrocytes and NIRG cells (*NKX2-2, OLIG2*), amacrine cells (*TFAP2A, PAX6*), bipolar cells (*OTX2*, *VSX2*), cone photoreceptors (*ISL2*, *CALB1*, *GNAT2*), and rod photoreceptors (*RHO*) (Fig. 9c). We probed for patterns and levels of chromatin access for *ID*s and *CDKN1A* in feature plots of all retinal cells (Fig. 9d-h). Similar to mRNA levels (Fig. 2), *ID2* chromatin accessibility was scattered across bipolar cells, amacrine cells, and photoreceptors (Fig. 9f). Unlike mRNA and protein levels (Fig. 2), *ID4* chromatin accessibility was apparent in some neurons as well as MG (Fig. 10h).

**Figure 9:**
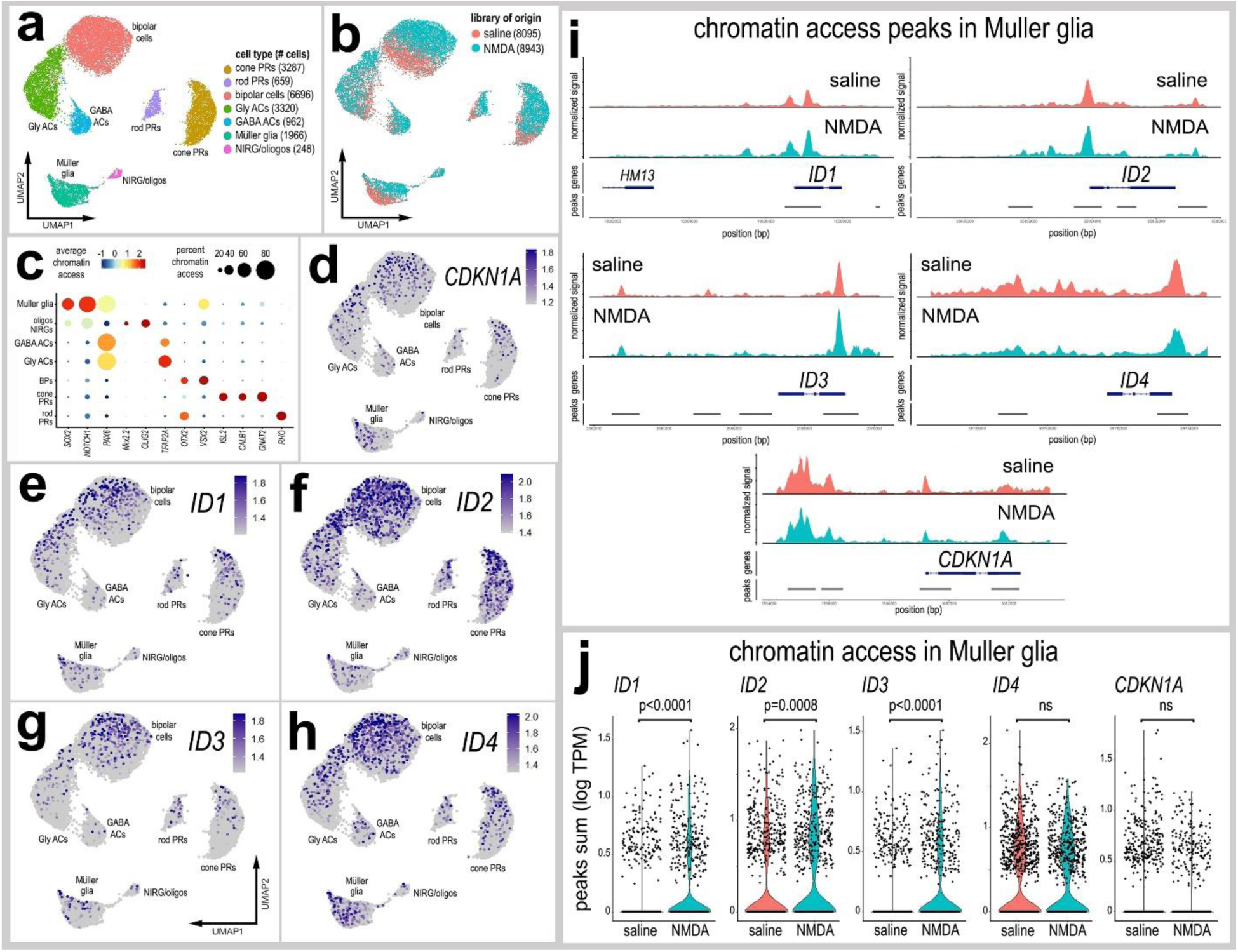
Chromatin access to ID4 and p21Cip1 in normal and damaged retinas. scATAC-seq was used to identify chromatin access of *ID*s and *CDKN1A* in control and NMDA-damaged retinas 24 hours after treatment. UMAP-ordered cells formed distinct clusters of neuronal cells, MG, NIRG cells and oligodendrocytes (**a,b**,**c**). Dot plot illustrates average accessibility (heatmap) and percent accessible (dot size) for different genes known to be associated with different retinal cell types (**c**). UMAP heatmap plots illustrate patterns and levels of access for *CDKN1A*, *ID1*, *ID2*, *ID3* and *ID4* across all retinal cell types (**d-h**). Coverage plots show chromatin access peaks across gene regions for *CDKN1A*, *ID1*, *ID2*, *ID3* and *ID4* in bioinformatically isolated MG from control and damaged retinas (**i**). Violin plots illustrate the levels for sums of chromatin access for *ID*s and *CDKN1A* in MG (**j**). Significance of difference (p-values) was determined by using a Wilcox rank sum test with Bonferroni correction.

**Figure 10:**
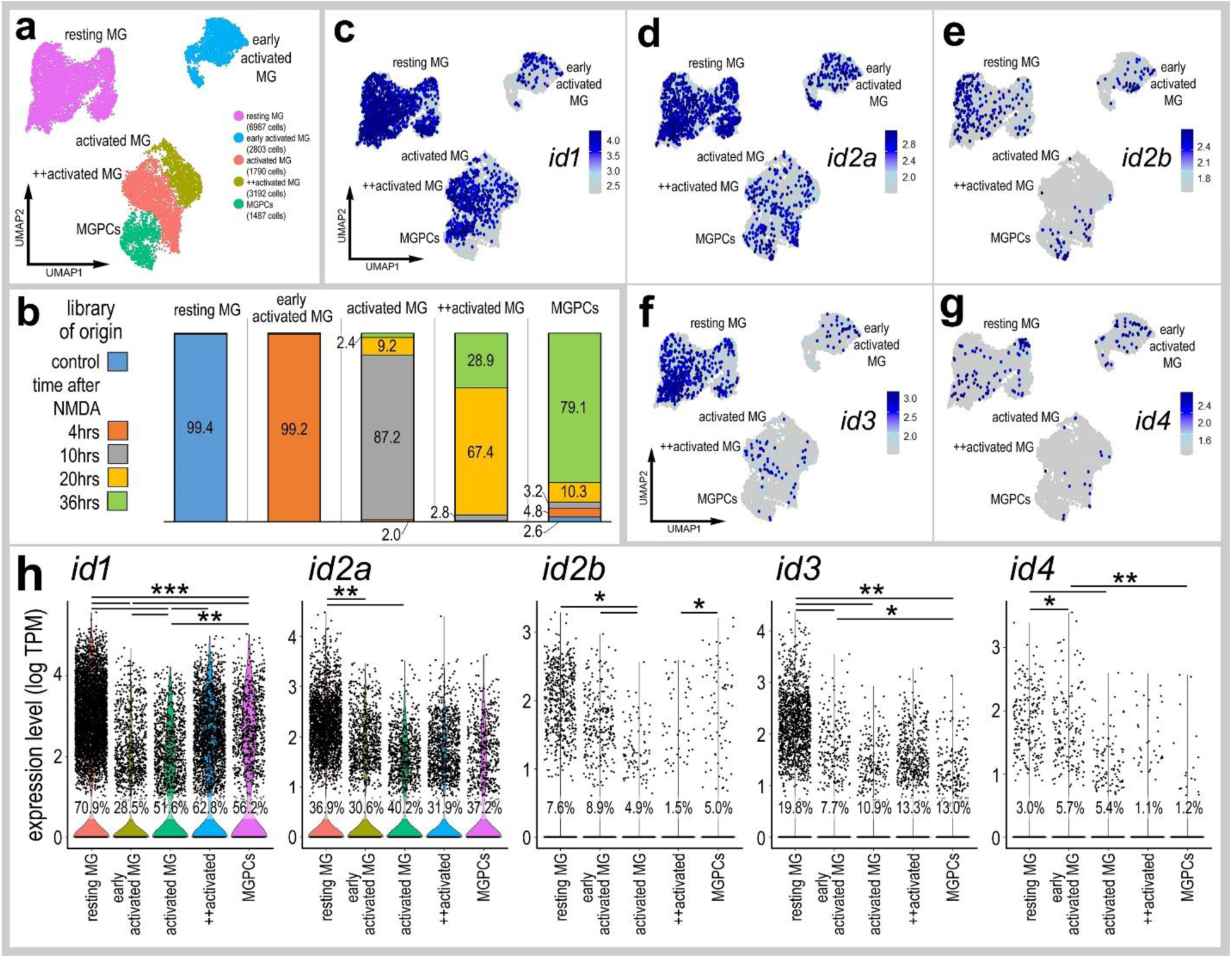
scRNA-seq for ID TFs in zebrafish MG. scRNA-seq was used to identify patterns and levels of expression of *id1, id2a, id2b, id3* and *id4* in control and NMDA-damaged retinas at 4, 10, 20 and 36 hours after treatment. UMAP clusters of cells were identified based on well-established patterns of gene expression (see Methods and supplemental Figure 1). Each dot represents one cell and black dots indicate cells that express 2 or more genes (**c-d**). The violin plots in **h** illustrate the expression levels and percent expressing cells *id1, id2a, id2b, id3* and *id4* in UMAP clusters of MG. Significant of difference (*p<0.01, **p<0.0001, ***p<<0.0001) was determined by using a Wilcox rank sum test with Bonferroni correction.

**Figure 11:**
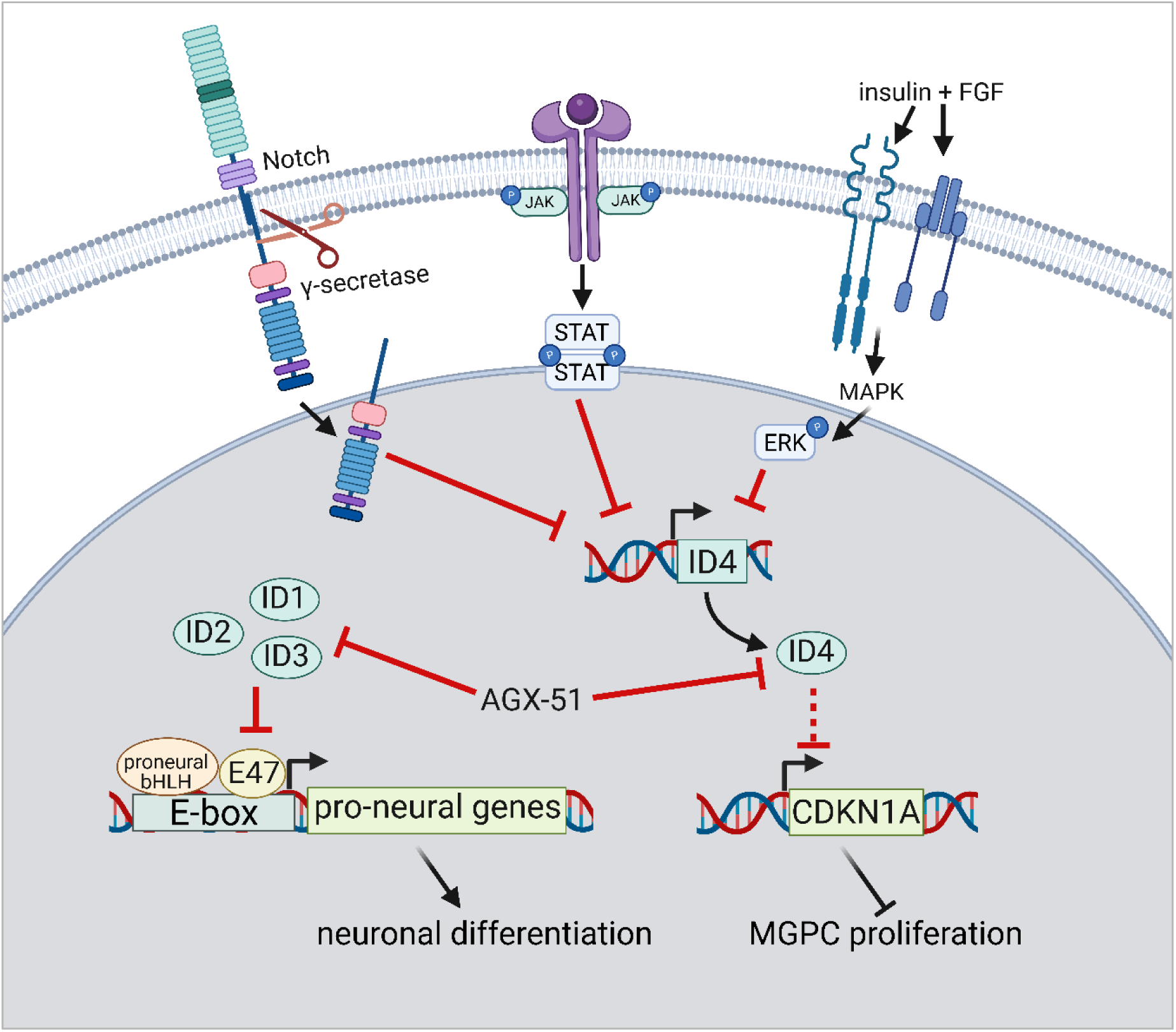
Schematic summary of findings. Notch-, gp130-, insulin- and FGF-signaling pathways are upstream regulators of *ID4* expression in MG. Applying ID inhibitor (AGX51) prior to neuronal damage, when levels of ID4 are high in MG, results in the upregulation of *CDKN1A* (p21^Cip1^) and suppressed proliferation of MGPCs. Inhibition of IDs after the formation of MGPCs, when levels of *ID1, ID2* and *ID3* are high in MGPCs, results in increased neuronal differentiation. This figure was generated using Biorender.com

We next probed for changes in chromatin access in MG in saline and NMDA-treated retinas. We assessed peak counts within the coding and flanking regions of *ID*s and *CDKN1A* (Fig. 9i). We identified significant increases in chromatin access in *ID1, ID2* and *ID3* in MG in NMDA-damaged retinas compared to access for these genes in resting MG (Fig. 9i,j). However, damage did not change chromatin access for *ID4* or *CDKN1A* in MG (Fig. 9i,j). Collectively, these findings suggest that chromatin access to *ID4* and C*DKN1A* is unaffected in MG in response to neuronal damage, whereas chromatin access to *ID1, ID2* and *ID3* is increase in MG in response to damage, which parallels mRNA levels.

### Patterns of ID expression in zebrafish retina

In undamaged mouse retinas, resting MG express very low levels of ID factors, but rapidly and transiently upregulate *Id1, Id2* and *Id3* is response to neuronal damage (Palazzo et al., 2022). This upregulation is, in part, downstream of NFkB signaling and inhibiting ID factors increases the differentiation of neuron-like cells from Ascl1-overexpressing MG (Palazzo et al., 2022). In the zebrafish retina, *Id2b* has been shown to be downregulated by MG in damaged retinas, and its overexpression suppresses the proliferation of MGPCs (Sahu et al., 2021). Midkine-A signaling is upstream of Id2a expression, which sustains the proliferation of progenitors in developing zebrafish retinas (Luo et al., 2012). Accordingly, we sought to better characterize patterns and levels of expression of ID factors in MG and MGPCs from scRNA-seq libraries of normal and NMDA-damaged zebrafish retinas, in previously described libraries (Hoang et al., 2020). MG and MGPCs for distinct clusters of UMAP-ordered cells and these clusters were largely populated by cells from different times after NMDA-treatment (Fig. 10a,b). Levels of *id1, id2a* and *id3* were relatively high in resting MG, decreased in early activated (4hrs after NMDA) MG and activated MG (predominantly occupied by MG from 10hrs after NMDA) (Fig. 10c,d,f,h). Levels of *id2a* and *id3* remained low in +activated MG (predominantly occupied by MG from 20hrs after NMDA) and MGPCs (predominantly occupied by MG from 36hrs after NMDA) (Fig. 10c,d,h). By comparison, levels of *id1* were increased in +activated MG and MGPCs compared to MG from earlier times after NMDA treatment (Fig. 10c,h). By comparison, *id2b* and *id4* were not highly expressed (>10% percent expression) by resting MG, activated MG or MGPCs (Fig. 10e,g,h). These data indicate that the downregulation of some *id* factors in MG after damage may be a necessary component of reprogramming in the zebrafish retina.

## Discussion

### Context-depended effects of IDs on proliferation, reprogramming and neuronal differentiation of MGPCs

We find dynamic patterns of expression of IDs in developing, normal and damaged chick retinas. These patterns of expression suggest that IDs have context-specific functions in MG. For example, levels of ID4 are high in resting MG, but levels are rapidly upregulated in response to neuronal damaged and downregulated by 12-24hrs after damage to remain low in MGPCs. By contrast, levels of ID1/2/3 are low in resting MG and upregulated in MGPCs. Application of ID inhibitor while levels of ID4 are high and transiently upregulated suppressed the formation of proliferating MGPCs in damaged retinas. Application of ID inhibitor while levels of ID1/2/3 are elevated resulted in increased neuronal differentiation of progeny produced by MGPCs in damaged retinas. These findings suggest that transient upregulation of ID4 is among the early steps that promote the reprogramming of MG into proliferating MGPCs, while elevated ID1/2/3 in MGPCs suppresses neuronal differentiation of progeny. Consistent with these observations, in the zebrafish retina, wherein neuronal differentiation from MGPC progeny is a robust process, the levels of IDs are greatly reduced (current study and (Sahu et al., 2021). Our findings suggest that inhibition of IDs promotes the increased differentiation of amacrine-like cells; this may reflect inherent specification-bias of MGPCs in the NMDA-damaged chick retina. We previously reported that inhibition of IDs enhanced regeneration of bipolar-like cells in NMDA-damaged Ascl1-overexpressing mouse retinas (Palazzo et al., 2022). However, the bias toward differentiation of bipolar-like cells is likely a function of elevated levels of Ascl1 imposing cell-type specification in mature MG. Thus, it seems likely that suppression of ID TFs enables increased neuronal differentiation without directing cell-type specification.

### IDs and cell survival

We observed decreased cell survival in damaged retinas treated with ID inhibitor. It is possible that decreased neuronal survival resulted from inhibition of ID factors in MG, which secondarily influences neuronal survival. Inhibition of different cell signaling pathways specifically in MG has been shown to have significant impacts on neuronal survival in damaged retinas (Gallina et al., 2015b; Todd et al., 2016b, 2017; Palazzo et al., 2020). Alternatively, since amacrine, bipolar and horizontal cells express ID2, it is possible that the ID inhibitor directly influenced the survival of these neurons in NMDA damaged retinas; we did not observe cell death in undamaged retinas treated with ID inhibitor alone (not shown). Some groups have shown that knockdown of ID2 drives excess mitochondrial activation and DNA damage in stem cells and glioblastoma cells (Zhang et al., 2017; Jakubison et al., 2022). Conversely, other groups have reported that ID2 is highly expressed in dying neurons and that ID2 knockdown prevents neuronal apoptosis (Gleichmann et al., 2002; Guo et al., 2015). Therefore, the cell death we observed in the inner INL of ID-inhibitor treated damaged retinas may have resulted from inhibition of ID2 function in amacrine cells. A third possibility is that the ID inhibitor had toxic off-target effects that negatively influenced the survival of neurons in damaged retinas. It is also possible that neuronal survival was diminished by ID inhibitor by a combination of all of the above-mentioned mechanisms.

### IDs in retinal development

In the developing chick retina, we found that expression levels of ID1, ID2 and ID4 peaked at E12 in RPCs and MG, and levels ID4 remained high in maturing MG. Previous reports have demonstrated the roles of IDs in zebrafish and mouse retinal development. In the fish, id2a loss-of-function drives premature cell cycle exit of retinal progenitors resulting in microphthalmia, over-production of RGCs, and a paucity of cell types normally found in INL and ONL (Uribe and Gross, 2010). During early murine ocular development, Id1, Id2, and Id3 expression is low in the optic vesicles and highly upregulated in the presumptive neural retina (Jen et al., 1997). Further, murine and human retinal progenitors express high levels of Id1 and Id3 which either decline as they progress toward a neural fate or are maintained in differentiating MG (Nelson et al., 2011; Hu et al., 2019). Deletion of Id1 and Id3 in the developing mouse retina causes microophthalmia and precocious differentiation of progenitors (Du and Yip, 2011). Moreover, forced expression of Id1 and Id3 at P0 enhances proliferation of retinal progenitor and MG-like cells (Mizeracka et al., 2013). Collectively, our data is consistent with the notion that IDs act to promote the proliferation of progenitors and bias specification toward a glial fate at the expense of neuronal fate.

### Pro-glial factors and the reprogramming of MG into MGPCs

Expression patterns of IDs in the developing retina mimic other pro-glial factors such as NFI’s and Notch-related factors. In the developing mouse retina, NFIs are required for cell cycle exit and the specification of bipolar cells and MG (Clark et al., 2019). (El-Hodiri et al., 2022). In the mature mouse retina, conditional knock-out of Nfia, Nfib and Nfix in mature MG combined with NMDA-induced damage results in the formation of MGPCs that produce some progeny that differentiate as amacrine- and bipolar-like neurons (Hoang et al., 2020). Similar to patterns of expression of ID4, NFIA, NFIB, NFIC and NFIX are expressed at relatively high levels in resting MG and are downregulated following NMDA-treatment and in MGPCs (El-Hodiri et al., 2022). Similar to other pro-glial factors, Notch-signaling and downstream bHLH factor Hes1 are required for the formation of MG during retinal development (Furukawa et al., 2000). In the fish retina, Notch-signaling must be downregulated in MG to initiate the process of regeneration (Conner et al., 2014; Elsaeidi et al., 2018; Campbell et al., 2021a; Sahu et al., 2021). Patterns of expression of NFIs and Ids suggest that these factors must be downregulated in MGPCs in damaged zebrafish retinas for neuronal regeneration to occur. Unlike patterns of expression in chick MGPCs, levels of id1, id2a and id3 are greatly decreased in zebrafish MGPCs (current study) and NFIs are not highly expressed by zebrafish MG (Hoang et al., 2020). The elevated levels of ID1/2/3 in chick MGPCs may, in part, explain why their neurogenic potential is very limited compared to that of zebrafish MGPCs. These observations are consistent with the notion that “pro-glial” factors significantly impact the resting phenotype of MG and their ability to reprogram into proliferating, neurogenic progenitor cells.

### Effects of ID inhibitor on MGPCs in retinas treated with insulin and FGF2

Although ID inhibitor suppressed the formation of proliferating MGPCs in NMDA-damaged retinas, this inhibitor had no effect upon the formation of MGPCs in undamaged retinas treated with insulin and FGF2. It is possible that ID inhibitor had no effect upon the proliferation of MGPCs in undamaged retinas because treatment with insulin and FGF2 potently downregulated ID4 leaving the inhibitor with nothing to act upon. By contrast, we found that NMDA-treatment rapidly upregulated ID4. The context, timing and levels of ID expression by MG in NMDA-damaged retinas is very different than in undamaged retinas treated with insulin and FGF2. The rapid, transient upregulation of ID4 may initiate a program that stimulates the proliferation of MGPCs in damaged retinas. Further, the percentage of expressing cells and levels of expression of ID1, ID2 and ID3 are higher in MG and MGPCs in NMDA-damaged retinas compared to levels seen in MG and MGPCs in retinas treated with insulin and FGF2 (see Fig. 2).

In general, most treatments have similar effects upon the proliferation of MGPCs in NMDA-damaged and insulin+FGF2-treated retinas. These treatments include activation or inhibition of MAPK (Fischer et al., 2009a; b), inhibition of Notch-signaling (Ghai et al., 2010), activation of glucocorticoid-signaling (Gallina et al., 2014b), activation or inhibition of Hedgehog-signaling (Todd and Fischer, 2015), activation or inhibition of Jak/Stat-signaling (Todd et al., 2016b), inhibition of mTor-signaling (Zelinka et al., 2016), activation or inhibition of retinoic acid-signaling (Todd et al., 2018), inhibition of BMP/Smad1/5/8-signaling (Todd et al., 2017), inhibition of Smad3-signaling (Todd et al., 2017), inhibition of MMP2/9 (Campbell et al., 2019), inhibition or activation of NFkB signaling (Palazzo et al., 2020), and inhibition of fatty acid binding proteins (Hoang et al., 2020; Campbell et al., 2022). Interestingly, activated microglia are required for the formation of proliferating MGPCs in damaged and undamaged retinas (Fischer et al., 2014b). By comparison, HB-EGF or EGFR-inhibitor (Todd et al., 2015) or activation or inhibition of midkine signaling (Campbell et al., 2021c) significantly influenced the formation of proliferating MGPCs in damaged retinas but not in retinas treated with FGF2 and insulin. Interestingly, we found that that the proliferation of MGPCs was unaffected by ID inhibitor; wherein levels of ID4 were potently downregulated by treatment with insulin and FGF2. Thus, damage-dependent pathways that regulate the reprogramming of MG into MGPCs may include distinct ID4-related pathways and contexts that are not active in undamaged retinas treated with insulin and FGF2.

### Changes in levels of ID4 and p21^Cip1^ in MG

We found that levels of ID4 in MG are paralleled, in most instances, by levels of p21^Cip1^. To the best of our knowledge, there are no reports that directly link ID4 to regulating expression levels of p21^Cip1^. Nevertheless, our findings suggest that some of the effects of ID inhibition may manifest, in part, by influencing levels of p21^Cip1^. Since p21^Cip1^ is a Cyclin-Dependent Kinase inhibitor, it is expected that diminished levels of p21^Cip1^ in MG correspond to increased proliferation of MGPCs. In undamaged retinas treated with insulin, FGF2 and ID inhibitor, we find that levels of p21^Cip1^ in MG were not increased and the ID inhibitor did not influence the proliferation of MGPCs. By contrast, in damaged retinas treated with ID inhibitor, levels of p21^Cip1^ in MG were increased and this treatment suppressed the proliferation of MGPCs. By comparison, damaged retinas treated with Notch or gp130 inhibitor increased levels of ID4 in MG, but did not influence levels of p21^Cip1^, wherein these inhibitors are known to suppress the proliferation of MGPCs (Hayes et al., 2007; Ghai et al., 2010; Todd et al., 2016b). These findings are consistent with the notion that levels of ID4 need to be upregulated, and then downregulated to “kick-start” the process of MG reprogramming into proliferation of MGPCs in damaged retinas. Further, these findings suggest that inhibition of Notch-or gp130-signaling does not suppress MGPC proliferation by upregulating p21^Cip1^.

## Conclusions

IDs are dynamically expressed in retinas following acute damage or treatment with growth factors that stimulate the formation of MGPCs. Dynamic changes in expression of mRNA strongly correlates with changes in protein function (Liu et al., 2016). Inhibition of ID TFs suppress the formation of proliferating MGPCs in damaged retinas, and this may occur, in part, by increasing levels of p21^Cip1^ in MG. Patterns of expression of p21^Cip1^ in MG parallel patterns of expression of ID4. Inhibition of IDs after proliferation increased the number of progeny that differentiate as amacrine-like cells. Our findings suggest that ID4 is highly expressed by maturing MG and resting MG and may act to maintain the phenotype resting MG, while a rapid upregulation after damage may drive the proliferation of MGPCs. By comparison, levels of ID1, ID2 and ID3 are increased in MGPCs, and these IDs may act to suppress neurogenic potential.

## Author Contributions

OT designed and executed experiments, gathered data, constructed figures and wrote the manuscript. SP and EH executed experiments and gathered data. HE and AJF designed experiments, analyzed data, constructed figures and wrote the manuscript.

## Data availability

Cell Ranger output files for Gene-Cell matrices for scRNA-seq data for libraries from saline and NMDA-treated retinas are available through GitHub: https://github.com/jiewwwang/Singlecell-retinal-regeneration or Sharepoint: [chick embryonic retina scRNA-seq Cell Ranger outs](https://osumc.sharepoint.com/:f:/s/Links/EtBjFrvATk9AuO_UBNr_LQABWnxJLKcLdfeVMTs3bPzjzw?e=FU3Sll) and [chick retina scRNA-seq Cell Ranger output files](https://osumc.sharepoint.com/:f:/s/Links/Eoto-Qg2uuxDn1bHWMM6gdkBTft4S_YSBjQJResxY-qehA?e=xMKHgS). scRNA-seq datasets are deposited in GEO (GSE135406, GSE242796) and Gene-Cell matrices for scRNA-seq data for libraries from saline and NMDA-treated retinas are available through NCBI (GSM7770646, GSM7770647, GSM7770648, GSM7770649).

---

**Supplemental Figure 1.**
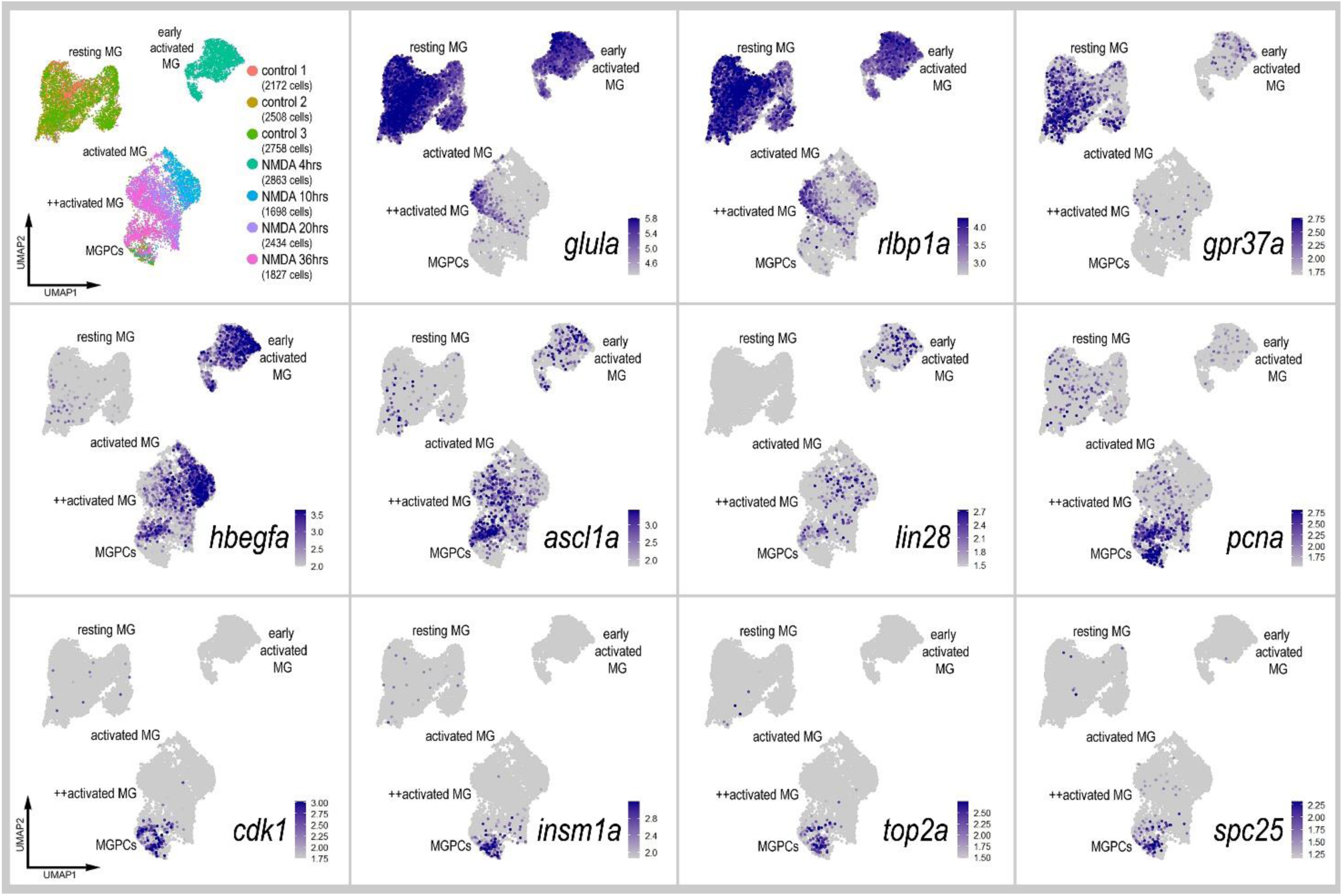
scRNA-seq for different progenitor- and MG-related genes in zebrafish MG. scRNA-seq was used to identify MG in control and NMDA-damaged retinas at 4, 10, 20 and 36 hours after treatment. UMAP clusters of cells were identified based on well-established patterns of gene expression, including *vim, glula, rlbp1a, gpr37a, hbegfa, ascl1a, lin28a, pcna, cdk1, insm1a, top2a* and *spc25*. Each dot represents one cell.

## References

Barbosa-Sabanero K, Hoffmann A, Judge C, Lightcap N, Tsonis PA, Del Rio-Tsonis K. 2012. Lens and retina regeneration: new perspectives from model organisms. Biochem J 447:321–34.

Bernardos RL, Barthel LK, Meyers JR, Raymond PA. 2007. Late-stage neuronal progenitors in the retina are radial Muller glia that function as retinal stem cells. J Neurosci 27:7028– 40.

Campbell LJ, Hobgood JS, Jia M, Boyd P, Hipp RI, Hyde DR. 2021a. Notch3 and DeltaB maintain Müller glia quiescence and act as negative regulators of regeneration in the light-damaged zebrafish retina. Glia 69:546–566.

Campbell WA, Blum S, Reske A, Hoang T, Blackshaw S, Fischer AJ. 2021b. Cannabinoid signaling promotes the de-differentiation and proliferation of Müller glia-derived progenitor cells. Glia 69:2503–2521.

Campbell WA, Deshmukh A, Blum S, Todd L, Mendonca N, Weist J, Zent J, Hoang TV, Blackshaw S, Leight J, Fischer AJ. 2019. Matrix-metalloproteinase expression and gelatinase activity in the avian retina and their influence on Müller glia proliferation. Exp Neurol 320:112984.

Campbell WA, El-Hodiri HM, Torres D, Hawthorn EC, Kelly LE, Volkov L, Akanonu D, Fischer AJ. 2023. Chromatin access regulates the formation of Müller glia-derived progenitor cells in the retina. Glia 71:1729–1754.

Campbell WA, Fritsch-Kelleher A, Palazzo I, Hoang T, Blackshaw S, Fischer AJ. 2021c. Midkine is neuroprotective and influences glial reactivity and the formation of Müller glia-derived progenitor cells in chick and mouse retinas. Glia 69:1515–1539.

Campbell WA, Tangeman A, El-Hodiri HM, Hawthorn EC, Hathoot M, Blum S, Hoang T, Blackshaw S, Fischer AJ. 2022. Fatty acid-binding proteins and fatty acid synthase influence glial reactivity and promote the formation of Müller glia-derived progenitor cells in the chick retina. Development 149:dev200127.

Carey JPW, Asirvatham AJ, Galm O, Ghogomu TA, Chaudhary J. 2009. Inhibitor of differentiation 4 (Id4) is a potential tumor suppressor in prostate cancer. BMC Cancer 9:173.

Clark BS, Stein-O’Brien GL, Shiau F, Cannon GH, Davis-Marcisak E, Sherman T, Santiago CP, Hoang TV, Rajaii F, James-Esposito RE, Gronostajski RM, Fertig EJ, Goff LA, Blackshaw S. 2019. Single-Cell RNA-Seq Analysis of Retinal Development Identifies NFI Factors as Regulating Mitotic Exit and Late-Born Cell Specification. Neuron 102:1111–1126.e5.

Conner C, Ackerman KM, Lahne M, Hobgood JS, Hyde DR. 2014. Repressing Notch Signaling and Expressing TNFalpha Are Sufficient to Mimic Retinal Regeneration by Inducing Muller Glial Proliferation to Generate Committed Progenitor Cells. J Neurosci 34:14403– 19.

Du Y, Yip HK. 2011. The expression and roles of inhibitor of DNA binding helix-loop-helix proteins in the developing and adult mouse retina. Neuroscience 175:367–379.

El-Hodiri HM, Campbell WA, Kelly LE, Hawthorn EC, Schwartz M, Jalligampala A, McCall MA, Meyer K, Fischer AJ. 2022. Nuclear Factor I in neurons, glia and during the formation of Müller glia-derived progenitor cells in avian, porcine and primate retinas. J Comp Neurol 530:1213–1230.

Elsaeidi F, Macpherson P, Mills EA, Jui J, Flannery JG, Goldman D. 2018. Notch suppression collaborates with Ascl1 and Lin28 to unleash a regenerative response in fish retina, but not in mice. J Neurosci [Internet]. Available from: https://www.ncbi.nlm.nih.gov/pubmed/29378863

Fausett BV, Goldman D. 2006. A role for alpha1 tubulin-expressing Muller glia in regeneration of the injured zebrafish retina. J Neurosci 26:6303–13.

Fischer AJ. 2005. Neural regeneration in the chick retina. Prog Retin Eye Res 24:161–82.

Fischer AJ, Bosse JL, El-Hodiri HM. 2014a. The ciliary marginal zone (CMZ) in development and regeneration of the vertebrate eye. Exp Eye Res [Internet] 116:199–204. Available from: http://www.ncbi.nlm.nih.gov/entrez/query.fcgi?cmd=Retrieve&db=PubMed&dopt=Citation&list_uids=24025744

Fischer AJ, Dierks BD, Reh TA. 2002. Exogenous growth factors induce the production of ganglion cells at the retinal margin. Development 129:2283–91.

Fischer AJ, Reh TA. 2001. Müller glia are a potential source of neural regeneration in the postnatal chicken retina. Nat Neurosci 4:247–252.

Fischer AJ, Ritchey ER, Scott MA, Wynne A. 2008. Bullwhip neurons in the retina regulate the size and shape of the eye. Dev Biol [Internet] 317:196–212. Available from: http://www.ncbi.nlm.nih.gov/entrez/query.fcgi?cmd=Retrieve&db=PubMed&dopt=Citation&list_uids=18358467

Fischer AJ, Scott MA, Ritchey ER, Sherwood P. 2009a. Mitogen-activated protein kinase-signaling regulates the ability of Müller glia to proliferate and protect retinal neurons against excitotoxicity. Glia 57:1538–1552.

Fischer AJ, Scott MA, Tuten W. 2009b. Mitogen-activated protein kinase-signaling stimulates Muller glia to proliferate in acutely damaged chicken retina. Glia 57:166–81.

Fischer AJ, Scott MA, Zelinka C, Sherwood P. 2010. A novel type of glial cell in the retina is stimulated by insulin-like growth factor 1 and may exacerbate damage to neurons and Muller glia. Glia 58:633–49.

Fischer AJ, Seltner RLP, Poon J, Stell WK. 1998. Immunocytochemical characterization of quisqualic acid- and N-methyl-D-aspartate-induced excitotoxicity in the retina of chicks. Journal of Comparative Neurology 393:1–15.

Fischer AJ, Zelinka C, Gallina D, Scott MA, Todd L. 2014b. Reactive microglia and macrophage facilitate the formation of Muller glia-derived retinal progenitors. Glia 62:1608–28.

Furukawa T, Mukherjee S, Bao ZZ, Morrow EM, Cepko CL. 2000. rax, Hes1, and notch1 promote the formation of Muller glia by postnatal retinal progenitor cells. Neuron 26:383– 94.

Gallina D, Palazzo I, Steffenson L, Todd L, Fischer AJ. 2015a. Wnt/betacatenin-signaling and the formation of Muller glia-derived progenitors in the chick retina. Dev Neurobiol [Internet]. Available from: http://www.ncbi.nlm.nih.gov/entrez/query.fcgi?cmd=Retrieve&db=PubMed&dopt=Citation&list_uids=26663639

Gallina D, Todd L, Fischer AJ. 2014a. A comparative analysis of Muller glia-mediated regeneration in the vertebrate retina. Exp Eye Res 123:121–130.

Gallina D, Zelinka C, Fischer AJ. 2014b. Glucocorticoid receptors in the retina, Muller glia and the formation of Muller glia-derived progenitors. Development 141:3340–51.

Gallina D, Zelinka CP, Cebulla CM, Fischer AJ. 2015b. Activation of glucocorticoid receptors in Müller glia is protective to retinal neurons and suppresses microglial reactivity. Exp Neurol 273:114–125.

Ghai K, Zelinka C, Fischer AJ. 2009. Serotonin released from amacrine neurons is scavenged and degraded in bipolar neurons in the retina. J Neurochem 111:1–14.

Ghai K, Zelinka C, Fischer AJ. 2010. Notch signaling influences neuroprotective and proliferative properties of mature Müller glia. J Neurosci 30:3101–3112.

Gleichmann M, Buchheim G, El-Bizri H, Yokota Y, Klockgether T, Kügler S, Bähr M, Weller M, Schulz JB. 2002. Identification of inhibitor-of-differentiation 2 (Id2) as a modulator of neuronal apoptosis. J Neurochem 80:755–762.

Guo L, Yang X, Lin X, Lin Y, Shen L, Nie Q, Ren L, Guo Q, Que S, Qiu Y. 2015. Silencing of Id2 attenuates hypoxia/ischemia-induced neuronal injury via inhibition of neuronal apoptosis. Behav Brain Res 292:528–536.

Hayes S, Nelson BR, Buckingham B, Reh TA. 2007. Notch signaling regulates regeneration in the avian retina. Dev Biol 312:300–11.

Hoang T, Wang J, Boyd P, Wang F, Santiago C, Jiang L, Yoo S, Lahne M, Todd LJ, Jia M, Saez C, Keuthan C, Palazzo I, Squires N, Campbell WA, Rajaii F, Parayil T, Trinh V, Kim DW, Wang G, Campbell LJ, Ash J, Fischer AJ, Hyde DR, Qian J, Blackshaw S. 2020. Gene regulatory networks controlling vertebrate retinal regeneration. Science 370:eabb8598.

Hu Y, Wang X, Hu B, Mao Y, Chen Y, Yan L, Yong J, Dong J, Wei Y, Wang W, Wen L, Qiao J, Tang F. 2019. Dissecting the transcriptome landscape of the human fetal neural retina and retinal pigment epithelium by single-cell RNA-seq analysis. PLoS Biol 17:e3000365.

Iavarone A, Lasorella A. 2004. Id proteins in neural cancer. Cancer Lett 204:189–196.

Jakubison BL, Sarkar T, Gudmundsson KO, Singh S, Sun L, Morris HM, Klarmann KD, Keller JR. 2022. ID2 and HIF-1α collaborate to protect quiescent hematopoietic stem cells from activation, differentiation, and exhaustion. J Clin Invest 132:e152599.

Jen Y, Manova K, Benezra R. 1997. Each member of the Id gene family exhibits a unique expression pattern in mouse gastrulation and neurogenesis. Dev Dyn 208:92–106.

Jorstad NL, Wilken MS, Grimes WN, Wohl SG, VandenBosch LS, Yoshimatsu T, Wong RO, Rieke F, Reh TA. 2017. Stimulation of functional neuronal regeneration from Muller glia in adult mice. Nature [Internet]. Available from: https://www.ncbi.nlm.nih.gov/pubmed/28746305

Jorstad NL, Wilken MS, Todd L, Finkbeiner C, Nakamura P, Radulovich N, Hooper MJ, Chitsazan A, Wilkerson BA, Rieke F, Reh TA. 2020. STAT Signaling Modifies Ascl1 Chromatin Binding and Limits Neural Regeneration from Muller Glia in Adult Mouse Retina. Cell Rep 30:2195–2208.e5.

Karl MO, Hayes S, Nelson BR, Tan K, Buckingham B, Reh TA. 2008. Stimulation of neural regeneration in the mouse retina. Proc Natl Acad Sci U S A 105:19508–13.

Knowell AE, Patel D, Morton DJ, Sharma P, Glymph S, Chaudhary J. 2013. Id4 dependent acetylation restores mutant-p53 transcriptional activity. Mol Cancer 12:161.

Liu Y, Beyer A, Aebersold R. 2016. On the Dependency of Cellular Protein Levels on mRNA Abundance. Cell 165:535–550.

Luo J, Uribe RA, Hayton S, Calinescu AA, Gross JM, Hitchcock PF. 2012. Midkine-A functions upstream of Id2a to regulate cell cycle kinetics in the developing vertebrate retina. Neural Dev 7:33.

Mizeracka K, DeMaso CR, Cepko CL. 2013. Notch1 is required in newly postmitotic cells to inhibit the rod photoreceptor fate. Development 140:3188–3197.

Murugesan P, Begum H, Tangutur AD. 2023. Inhibitor of DNA binding/differentiation proteins as IDs for pancreatic cancer: Role in pancreatic cancer initiation, development and prognosis. Gene 853:147092.

Nelson BR, Ueki Y, Reardon S, Karl MO, Georgi S, Hartman BH, Lamba DA, Reh TA. 2011. Genome-wide analysis of Muller glial differentiation reveals a requirement for Notch signaling in postmitotic cells to maintain the glial fate. PLoS One 6:e22817.

Palazzo I, Deistler K, Hoang TV, Blackshaw S, Fischer AJ. 2020. NF-κB signaling regulates the formation of proliferating Müller glia-derived progenitor cells in the avian retina. Development 147:dev183418.

Palazzo I, Todd LJ, Hoang TV, Reh TA, Blackshaw S, Fischer AJ. 2022. NFkB-signaling promotes glial reactivity and suppresses Müller glia-mediated neuron regeneration in the mammalian retina. Glia 70:1380–1401.

Powers AN, Satija R. 2015. Single-cell analysis reveals key roles for Bcl11a in regulating stem cell fate decisions. Genome Biol [Internet] 16:199. Available from: https://www.ncbi.nlm.nih.gov/pubmed/26390863

Ross SE, Greenberg ME, Stiles CD. 2003. Basic helix-loop-helix factors in cortical development. Neuron 39:13–25.

Sahu A, Devi S, Jui J, Goldman D. 2021. Notch signaling via Hey1 and Id2b regulates Müller glia’s regenerative response to retinal injury. Glia 69:2882–2898.

Satija R, Farrell JA, Gennert D, Schier AF, Regev A. 2015. Spatial reconstruction of single-cell gene expression data. Nat Biotechnol [Internet] 33:495–502. Available from: https://www.ncbi.nlm.nih.gov/pubmed/25867923

Sharma P, Patel D, Chaudhary J. 2012. Id1 and Id3 expression is associated with increasing grade of prostate cancer: Id3 preferentially regulates CDKN1B. Cancer Med 1:187–197.

Stuart T, Srivastava A, Madad S, Lareau CA, Satija R. 2021. Single-cell chromatin state analysis with Signac. Nat Methods 18:1333–1341.

Todd L, Finkbeiner C, Wong CK, Hooper MJ, Reh TA. 2020. Microglia Suppress Ascl1-Induced Retinal Regeneration in Mice. Cell Rep 33:108507.

Todd L, Fischer AJ. 2015. Hedgehog-signaling stimulates the formation of proliferating Müller glia-derived progenitor cells in the retina. Development 142:2610–2622.

Todd L, Hooper MJ, Haugan AK, Finkbeiner C, Jorstad N, Radulovich N, Wong CK, Donaldson PC, Jenkins W, Chen Q, Rieke F, Reh TA. 2021. Efficient stimulation of retinal regeneration from Müller glia in adult mice using combinations of proneural bHLH transcription factors. Cell Rep 37:109857.

Todd L, Palazzo I, Squires N, Mendonca N, Fischer AJ. 2017. BMP- and TGFbeta-signaling regulate the formation of Muller glia-derived progenitor cells in the avian retina. Glia [Internet]. Available from: https://www.ncbi.nlm.nih.gov/pubmed/28703293

Todd L, Palazzo I, Suarez L, Liu X, Volkov L, Hoang TV, Campbell WA, Blackshaw S, Quan N, Fischer AJ. 2019. Reactive microglia and IL1β/IL-1R1-signaling mediate neuroprotection in excitotoxin-damaged mouse retina. J Neuroinflammation 16:118.

Todd L, Reh TA. 2022. Comparative Biology of Vertebrate Retinal Regeneration: Restoration of Vision through Cellular Reprogramming. Cold Spring Harb Perspect Biol 14:a040816.

Todd L, Squires N, Suarez L, Fischer AJ. 2016a. Jak/Stat signaling regulates the proliferation and neurogenic potential of Müller glia-derived progenitor cells in the avian retina. Sci Rep 6:35703.

Todd L, Squires N, Suarez L, Fischer AJ. 2016b. Jak/Stat signaling regulates the proliferation and neurogenic potential of Muller glia-derived progenitor cells in the avian retina. Sci Rep 6:35703.

Todd L, Suarez L, Quinn C, Fischer AJ. 2018. Retinoic Acid-Signaling Regulates the Proliferative and Neurogenic Capacity of Müller Glia-Derived Progenitor Cells in the Avian Retina. Stem Cells 36:392–405.

Todd L, Volkov LI, Zelinka C, Squires N, Fischer AJ. 2015. Heparin-binding EGF-like growth factor (HB-EGF) stimulates the proliferation of Muller glia-derived progenitor cells in avian and murine retinas. Mol Cell Neurosci 69:54–64.

Torres-Machorro AL. 2021. Homodimeric and Heterodimeric Interactions among Vertebrate Basic Helix-Loop-Helix Transcription Factors. Int J Mol Sci 22:12855.

Uribe RA, Gross JM. 2010. Id2a influences neuron and glia formation in the zebrafish retina by modulating retinoblast cell cycle kinetics. Development 137:3763–3774.

Wan J, Goldman D. 2016. Retina regeneration in zebrafish. Curr Opin Genet Dev 40:41–47.

Wojnarowicz PM, Escolano MG, Huang Y-H, Desai B, Chin Y, Shah R, Xu S, Yadav S, Yaklichkin S, Ouerfelli O, Soni RK, Philip J, Montrose DC, Healey JH, Rajasekhar VK, Garland WA, Ratiu J, Zhuang Y, Norton L, Rosen N, Hendrickson RC, Zhou XK, Iavarone A, Massague J, Dannenberg AJ, Lasorella A, Benezra R. 2021. Anti-tumor effects of an ID antagonist with no observed acquired resistance. NPJ Breast Cancer 7:58.

Zelinka CP, Volkov L, Goodman ZA, Todd L, Palazzo I, Bishop WA, Fischer AJ. 2016. mTor signaling is required for the formation of proliferating Muller glia-derived progenitor cells in the chick retina. Development 143:1859–73.

Zhang Z, Rahme GJ, Chatterjee PD, Havrda MC, Israel MA. 2017. ID2 promotes survival of glioblastoma cells during metabolic stress by regulating mitochondrial function. Cell Death Dis 8:e2615.

